# A two-bead-per-aminoacid coarse-grained MD model with hydrogen bonding (2BPA-HB) to probe DNAJB6b-mediated suppression of polyglutamine aggregation in Huntington’s disease

**DOI:** 10.64898/2026.08.24.746793

**Authors:** Vasista Adupa, Johan D. Polet, Maurice Dekker, Patrick R. Onck

## Abstract

Polyglutamine (polyQ) aggregation plays a central role in several neurodegenerative diseases, including Huntington’s disease. DNAJB6b, a molecular chaperone involved in protein quality control, is known to efficiently suppress polyQ aggregation, but its anti-aggregation mechanism remains unclear. In this work we investigate the interaction between DNAJB6b and the polyQ region (Q48) of mutant Huntingtin Exon 1 (mHttEx1) using a custom-built coarse-grained molecular dynamics model. The model incorporates a two-bead-per-amino-acid representation with hydrogen bonding (termed 2BPA-HB), and is calibrated against all-atom molecular dynamics data in terms of geometry, hydrophobicity, and hydrogen bonding. The model reproduces the tertiary structure of DNAJB6b and its interactions with Q48, and reveals an inverse correlation between DNAJB6b concentration and Q48 aggregation propensity. Our simulations show that DNAJB6b co-condensates with polyQ molecules, thereby shielding the polyQ from forming the intermolecular hydrogen bonds necessary for amyloid formation. The 2BPA-HB CGMD model enables efficient exploration of DNAJB6b conformations, supporting future studies of chaperone-mediated aggregation suppression and therapeutic development.

## Introduction

Huntington’s disease (HD) is a neurodegenerative disorder characterized by a decline in motor control, behavioral and cognitive abilities^1^. The Huntingtin (Htt) protein is expressed in the cytoplasm of all human cells, with especially high levels in the brain. It is encoded by 67 exons and plays essential roles in embryonic development^2,3^, axonal vesicle transport^4^, and potentially in long-term memory storage^5^. The first exon of Htt (HttEx1) is well studied and consists of an N-terminal domain of 17 residues (N17), a stretch of glutamine (polyQ) residues, and a proline-rich domain (PRD) comprising 49 residues enriched in proline (P). In its non-pathogenic form, the polyQ tract in HttEx1 typically consists of 17–20 glutamines^6^; However, in HD, fragments of expanded polyQ (≥ 35) Htt are generated from protease activity and aberrant splicing. These fragments are prone to enhanced aggregation leading to the formation of fibrils and ultimately cellular inclusions, which are found as cytoplasmic and nuclear aggregates within neurons of the striatum, and is a well established hallmark of HD pathology^7–11^. Importantly, the age of onset is inversely correlated to the polyglutamine repeat length^12–15^.

Two primary mechanisms have been proposed to explain the formation of insoluble amyloid fibrils from monomeric species. The first is the nucleation elongation mechanism, in which a nucleus forms either by the association of more than four short monomers into an ordered oligomer, or when a single extended monomer folds into a β-hairpin structure^16^. This nucleus then serves as a seed for additional monomers to attach to its surface, thereby promoting fibril growth^17,18^. The second mechanism proposes that polyQ monomers undergo liquid–liquid phase separation, forming dense, liquid-like droplets, which undergo a liquid-to-solid phase transition, giving rise to β-sheet-rich structures^19–22^. These aggregates are toxic because they interfere with essential cellular components, such as proteins, membranes, and the proteostasis network—thereby disrupting cellular function^23^. Fortunately, cells are equipped with a protein quality control (PQC) system that acts as a defence mechanism against protein misfolding.

Molecular chaperones are central to protein quality control (PQC) network, supporting proteins throughout their life cycle from folding to assembly and degradation^24^. Among molecular chaperones, heat shock proteins (Hsps) are ubiquitous and conserved protein families in both prokaryotic and eukaryotic organisms^25^. One major system within this family is the 70 kDa Hsp (Hsp70), which are implicated in a variety of cellular process. Hsp70s never function alone; they are assisted by J-domain proteins (JDPs) and nucleotide exchange factors^26^. JDPs are the client recognition proteins, which then transfer the client to Hsp70 for further processing^26^.

One such JDP is DNAJB6b, a class B co-chaperone that stabilizes its client proteins in conformations compatible with proper folding or degradation. Preserving their disordered, liquid-like phase state of these clients is critical, as their transition into ordered, solid amyloid fibrils leads to toxicity and disease. DNAJB6b has been identified as a rate-limiting chaperone in suppressing polyglutamine aggregation, toxicity, and disease progression^27–31^. It is implicated in preventing protein aggregation across a range of disease-related and physiological conditions^32–40^.

In this study, we focus on DNAJB6b and its role in modulating polyglutamine aggregation associated with Huntington’s disease. DNAJB6b is the isoform b of DNAJB6 and is widely expressed in the brain^28^. It comprises four domains: the J-domain, which is a conserved feature of all JDPs; a glycine/phenylalanine-rich (G/F) region; a serine/threonine-rich (S/T) region; and a C-terminal domain (CTD). Structural studies have shown that the Hsp70-binding sites located on helices II and III are occluded by helix V from the G/F region, resulting in an autoinhibited conformation^41^. Our recent all-atom MD simulations demonstrated that DNAJB6b can sample three distinct conformations: (1) a closed state, (2) an open state, and (3) an extended state^42^. Importantly, the autoinhibitory state of DNAJB6b remains stable despite these conformational transitions. The precise molecular interactions by which this chaperone prevents the aggregation of disease-related proteins, however, remain poorly understood. Here, we aim to elucidate the molecular basis of DNAJB6b’s anti-aggregation mechanism against polyglutamine peptides using molecular dynamics (MD) simulations.

All-atom MD simulations provide atomistic detail of protein behavior, offering valuable insights into phenomena such as folding and aggregation. Despite their strengths, the high computational cost of all-atom MD restricts their applicability to small systems or short timescales—limitations that hinder the study of large-scale protein aggregation^43^. To overcome this, coarse-grained molecular dynamics (CGMD) models have been introduced, wherein multiple atoms are represented by a single interaction site. This abstraction drastically reduces computational cost, allowing for the simulation of larger systems over extended timescales, while preserving the key physicochemical features relevant to aggregation.

Coarse-grained protein models span a range of resolutions, from one-bead-per-residue approaches to representations containing multiple backbone and side-chain interaction sites^44–46^. One-bead models are computationally efficient and have been used extensively to investigate intrinsically disordered proteins and biomolecular phase separation^47–51^. Models containing additional interaction sites can provide a more detailed description of local geometry, side-chain specificity, and hydrogen-bond directionality, but generally require more extensive interaction potentials and parameterization. Examples include AWSEM and its IDP-specific variant AWSEM-IDP, which represent each residue using *C*_α_, *C*_β_, and backbone oxygen sites and combine physically motivated interactions with statistical and energy-landscape-based potentials^52,53^. Other multiscale coarse-grained models uses a *C*_α_ backbone site together with multiple side-chain sites, with its bonded and nonbonded interactions parameterized using atomistic simulations and structural distributions^54^. Similarly, UNRES, which forms the protein representation used within the UNICORN framework, describes each residue a using united peptide-group and side-chain interaction sites and employs an effective energy function containing local, nonlocal, and multibody contributions^55^.

The present 2BPA-HB model occupies a simpler position within this hierarchy. It contains one massive *C*_α_ backbone bead per residue together with amino-acid-specific virtual side-chain sites and virtual hydrogen-bond donor and acceptor sites. Only the equations of motion of the *C*_α_ backbone beads are integrated, whereas the positions of the virtual sites are calculated from the coordinates of the neighbouring backbone beads at each simulation step. The virtual-site parameters can therefore be obtained directly by mapping all-atom trajectories onto the corresponding coarse-grained geometry, without introducing additional independently integrated particles. This representation adds side-chain specificity and hydrogen-bonding while retaining one integrated particle per residue, allowing relatively large multichain systems to be simulated over the timescales required to observe polyQ aggregation.

A recent study by Dekker *et al*. introduced a CGMD model (2BPA-Q) that effectively captures the characteristic aggregation behavior of polyQ sequences^56^, driven by hydrogen bonding and beta-sheet formation. In the current work we extend this force field to account for all amino-acids (not just glutamine) for both the disordered and structured regions of DNAJB6b, and termed it 2BPA-HB, which is developed specifically for the DNAJB6b–Q48 system.

To achieve this, we calibrated the model through all-atom MD simulations of polyQ in complex with DNAJB6b. We first describe the development of the CGMD force field, which involved stepwise calibration of hydrophobic interactions, hydrogen bonding, and side chain parameters using data from all-atom MD simulations. We then apply the fully parameterized model to investigate how the molecular chaperone DNAJB6b influences polyglutamine aggregation. Initial single-molecule simulations of DNAJB6b confirmed that the model indeed captures DNAJB6b’s conformational transitions between open and extended states, consistent with all-atom MD data. In control simulations lacking the chaperone, polyQ molecules in solution form disordered aggregates, which eventually transition into ordered structures (i.e., amyloid like structures). Upon introducing DNAJB6b to a solution of polyQ molecules, we observe that the chaperone co-condensates with polyQ48 molecules inhibiting aggregation by preventing the formation of inter-polyQ hydrogen bonds that are critical for amyloid formation. When DNAJB6b is introduced after amyloid formation has occurred, however it is unable to disrupt the existing aggregates.

The 2BPA-HB model presented here provides a robust framework for future investigations into the molecular mechanisms of chaperone-mediated aggregation supression and therapeutic approaches for Huntington’s disease.

## Methods

The 2BPA-HB model extends a previously established one-bead-per-amino-acid (1BPA) framework, in which each residue is represented by a single coarse-grained bead centered at the C_*α*_ position. The 1BPA model includes backbone interactions appropriate for intrinsically disordered proteins (IDPs) and has been applied to study nucleocytoplasmic transport through the nuclear pore complex^44,57–59^ and liquid–liquid phase separation of transcription factors^60^, RNA-binding proteins^47,51^ and FG-Nups^50^. However, to accurately capture polyQ aggregation and its interaction with folded proteins like DNAJB6b, a more detailed CGMD model is required.

We therefore extended the 1BPA model to a 2BPA model that also accounts for a second (virtual) side chain bead (Fig. 1). Using virtual interaction sites for the side chains increases computational efficiency, with a minimal reduction of accuracy. Furthermore, two virtual interaction sites are used to capture hydrogen bonding interactions. The model is based on the 2BPA-Q model by Dekker et al.^56^ for polyQ and extended to include all amino-acid types. Essential differences in implementation will be highlighted.

**Figure 1:**
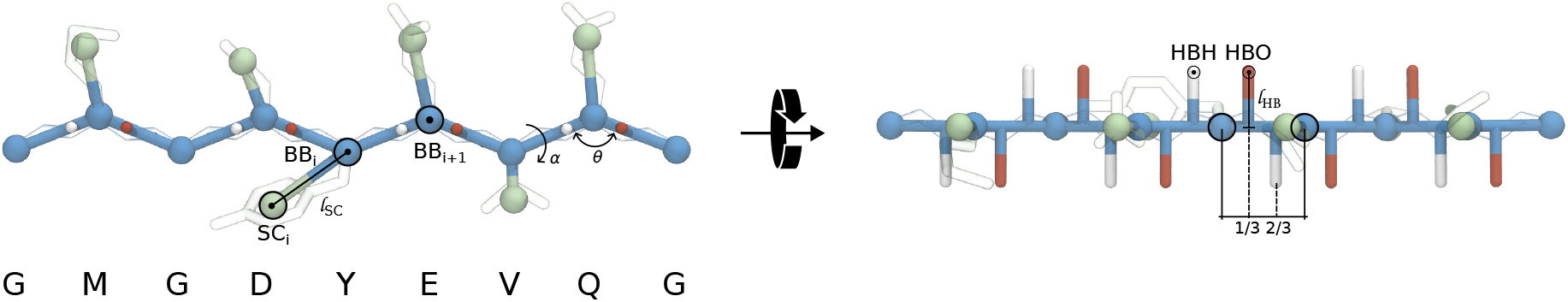
Geometry and bead representation of the 2BPA-HB model. Coarse-grained representation of a peptide chain composed of nine backbone beads. The backbone stiffness is modeled via pseudo-bending and torsion angles between the BB beads (blue). The various side chains are represented by virtual beads (SC_*i*_), whose positions are defined using the 3fd scheme with parameters *a* and *l*_SC_, calibrated specifically for each aminio acid as described in the Methods section. Hydrogen bonding beads for oxygen (HBO) and hydrogen (HBH) are placed using the 3out virtual site scheme, with parameters taken directly from Dekker *et al*.^56^. The hydrogen bond length (*l*_HB_ = 2.38 Å) corresponds to an interstrand distance of 4.76 Å in β-sheet structures. Both hydrogen bonding beads are positioned directly above and below the bond connecting BB_*i*_ and BB_*i*+1_. The coarse-grained structure is overlaid with the corresponding all-atom configuration (heavy atoms only, shown in transparency), illustrate the alignment between the all-atom side chain center of mass and the coarse-grained SC bead.

The bonded potential used for the backbone beads is adopted directly from the 1BPA model^44^ and is given by

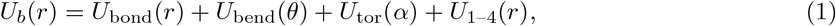

where *U*_bond_(*r*) is the distance dependent bond stretching term, *U*_bend_(*θ*) is the angle dependent bending term, *U*_tor_(*α*) is the angle dependent torsion term and *U*_1–4_(*r*) is the bending-torsion coupling term. The bond stretching is prescribed by a stiff harmonic potential with an equilibrium distance between two beads of 3.8 Å, and a stiffness of 8038 kJ nm^−2^ mol^−1^. The bending and torsion terms are both obtained from Ramachandran data and the Boltzmann inversion method^57^. The bending-torsion coupling potential ensure proper sampling in (*θ, α*)-space because of the uncoupled backbone dihedrals. For a detailed description of the potentials in Eq. 1, the reader is referred to Ghavami et al.^57^.

The non-bonded potential consists of two components: one accounting for hydrophobic and volume exclusion interactions (*U*_hp_(*r*)), and the other for electrostatics (*U*_el_(*r*)):

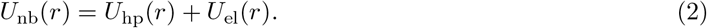

The hydrophobic potential *U*_hp_(*r*) incorporates both attractive and repulsive contributions combined into a single potential,

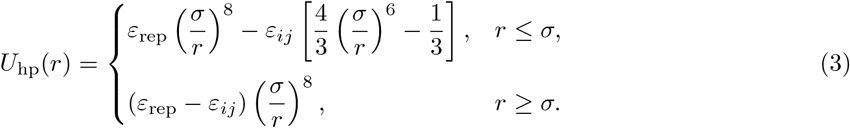

In this expression, *ε*_rep_ sets the strength of the repulsive force, while *σ* denotes the bead diameter, fixed at 4.76 Å for both backbone and side chain beads, following prior studies^50^, and *ε*_*ij*_ is the interaction strength between residues of type *i* and *j*. This interaction strength between pairs of residues can be determined by

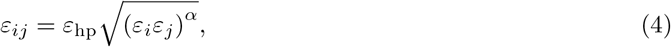

where *ε*_hp_ is the attractive hydrophobic interaction strength, *α* is a hydrophobic scaling factor which has to be fine-tuned, and *ε*_*i*_ is the hydrophobic strength of a bead of type *i*. The variables *ε*_hp_ = 13 kJ mol^−1^, and *ε*_rep_ = 10 kJ mol^−1^ are taken from the 1BPA model^44^. The hydrophobicity parameters *ε*_*i*_, are taken directly from Dekker *et al*. 2023^50^ (version 1BPA-1.1), listed in Supplementary Table 5.

The second term of the non-bonded potential is *U*_el_(*r*), and is given by

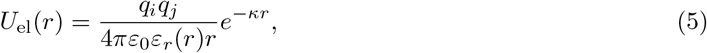

where *q*_*i*_ is the charge of residue *i, ε*_0_ is the dielectric vacuum permittivity, *κ* is the Debye screening coefficient, chosen to be 1.27 nm^−1^, and *ε*_*r*_(*r*) is the dimensionless distance-dependent dielectric coefficient given by the sigmoidal function

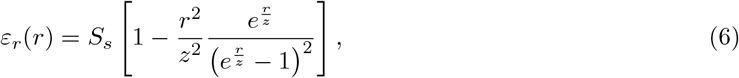

where the constants *z* = 2.5 Å and *S*_*s*_ = 80 are chosen^44,61^.

The 2BPA-HB model accounts for the side chain through a virtual site (in addition to the backbone bead) and captures hydrogen bonding through two virtual interactions sites per residue at the backbone (see Fig. 1). The hydrogen bonding sites mimic the spatial locations of the backbone NH (donor) and CO (acceptor) groups. Proline and glycine are treated as special cases: glycine lacks a side chain, and proline’s side chain forms a ring with the backbone nitrogen, preventing the assignment of a hydrogen bonding bead (HBH) in this model. The side chain bead positions follow the 3fd scheme, while the hydrogen bonding beads (HBO, HBH) follow the 3out scheme^62^ (see Fig. 1 and the Supplementary Information (SI), including Supplementary Fig. 3).

The interaction between the hydrogen bonding sites (a hydrogen and an oxygen bead) in the model is described by a shifted Lennard-Jones potential similar to the CGMD model of Chen and Imamura^63,64^,

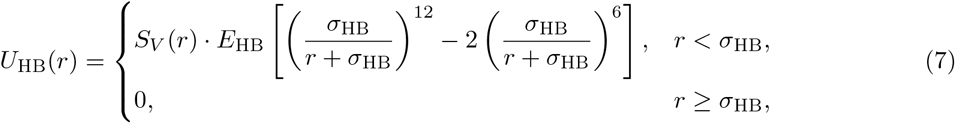

where *E*_HB_ is the hydrogen bonding energy, *σ*_HB_ = 5.12 Å is the range of the hydrogen bonding interaction, *r* is the distance between the H and O beads, and the scaling factor is defined as

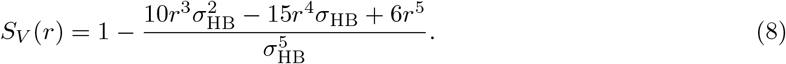

This potential is chosen such that the minimum of *U*_HB_(0) = −*E*_HB_ lies at *r* = 0. Furthermore, the scaling factor *S*_*V*_ (*r*) is introduced to ensure that the hydrogen bonding interaction has completely vanished at *r* = *σ*_HB_. Interactions between two H-beads or between two O-beads, respectively, are ignored in this model.

The total potential energy of the 2BPA-HB model is given by

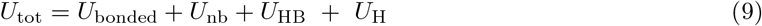

*U*_H_ represents the potential energy contribution from the elastic network, i.e., additional harmonic bonds used to preserve selected structural features of DNAJB6b (see Supplementary Eq. 1).

Our recent all-atom MD study revealed that DNAJB6b is a predominantly structured protein that interconverts between three conformational states: closed, open, and extended^42^. To represent the structured nature of DNAJB6b in the CGMD model while preserving its conformational flexibility, we applied elastic network restraints at two distinct levels. A fully connected elastic network with a harmonic bond constant of 8000 kJ mol^−1^ nm^−2^ was used to maintain the secondary structure of the protein, based on residue-level information from all-atom MD simulations (Fig. 2a). Details on the elastic network is provided in section *Elastic network* of the SI. In addition, a minimal set of extra elastic restraints was introduced to stabilize the overall tertiary structure while still allowing the protein to transition between different conformational states, particularly between the open and extended states (see supplementary section *Elastic network*, Supplementary Tab. 4). As a result, the final CGMD model successfully captures the interconversion between open and extended states, although transitions involving the closed state were not captured.

**Figure 2:**
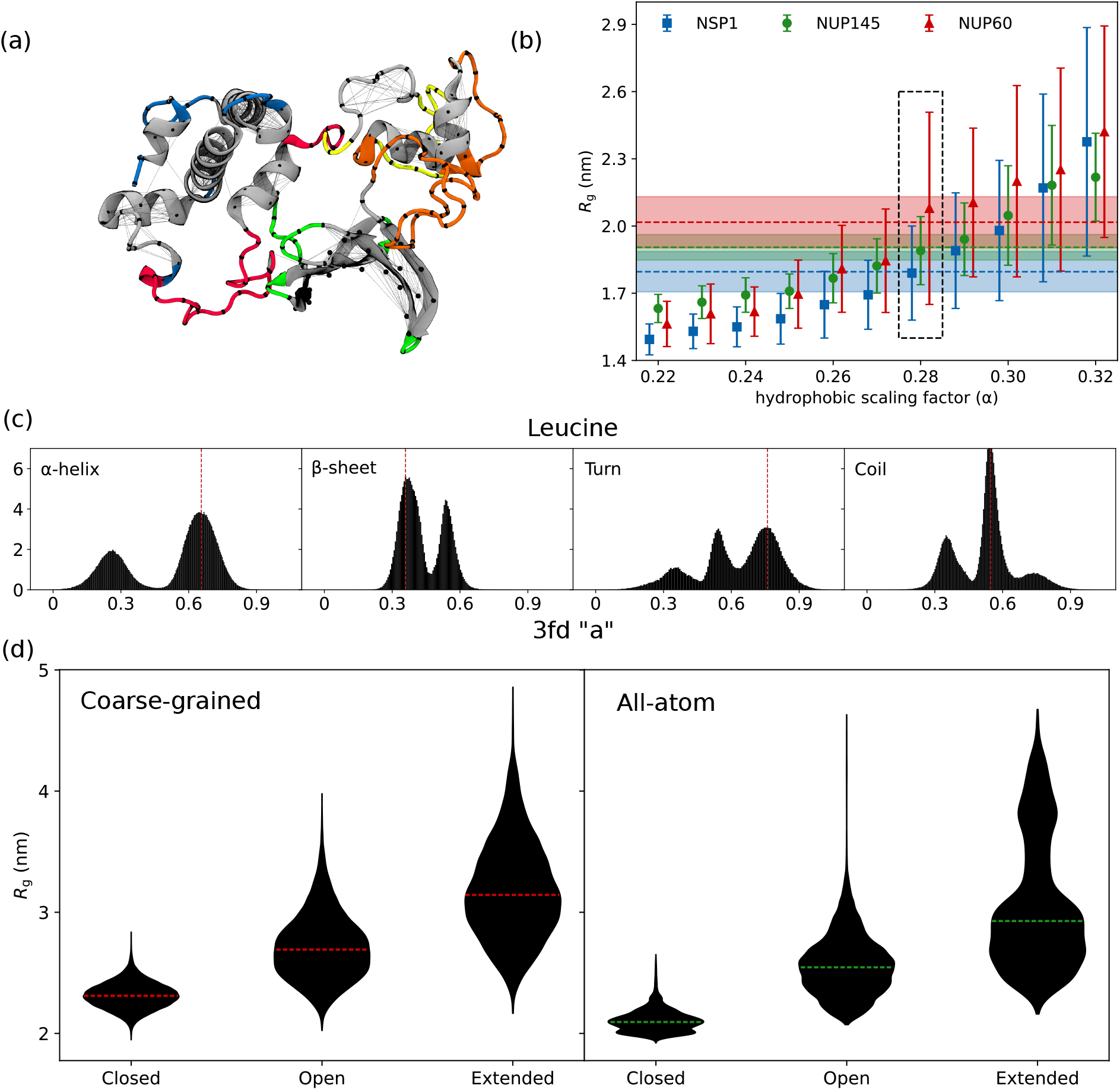
Calibration of parameters for DNAJB6b. **(a)** Snapshots showing the closed state of DNAJB6b with elastic network applied to retain the secondary structure of the protein. Each black line denotes a harmonic bond connecting two backbone beads, shown as black spheres. Colored regions are flexible, while grey regions are constrained using the elastic network. **(b)** Plots of *R*_g_ values from CGMD simulations of three FG-Nup segments as a function of the hydrophobic scaling factor (α_SC_), where error bars indicate the standard deviation observed across three repeats of the simulation. All-atom *R*_g_ values from three replicas are indicated by the shaded color regions for each FG-Nup and horizontal dashed lines represent the mean; the dashed black rectangle highlights the optimal α_SC_. **(c)** Probability density distributions of the `3fd` parameter *a* for leucine (as an example), derived from all-atom simulations of a single DNAJB6b molecule^42^. Distributions are separated by secondary structure type, showing that `3fd` parameter *a* strongly depends on local secondary structure. In contrast, the side chain length parameter |*b*| shows minimal variation across structures (Supplementary Tab. 2, Supplementary Fig. 4-21). Using the calibrated geometry and hydrophobicity parameters, CGMD simulations were performed to evaluate *R*_g_ for three states of DNAJB6b and plotted as violins **(d)**, where the mean of each plot is given by a horizontal dashed line, and compared to the all-atom data of a single molecule DNAJB6b^42^.

### Calibration

The calibration of the 2BPA-HB model was carried out in a stepwise manner, beginning with the optimization of parameters for DNAJB6b. This involved three main components: the geometry of the side chains, i.e., side chain virtual bead placements, the calibration of hydrophobicity and the evaluation of the secondary and tertiary structure. Once DNAJB6b was calibrated, the model parameters for polyglutamine (Q48) were optimized consisting of geometry calibration, and tuning the hydrophobicity parameter for the glutamine residues, after which the hydrogen bonding energy was calibrated to capture the amyloid formation. Lastly, the interaction parameters between DNAJB6b and polyQ were refined to reproduce the co-condensation behavior observed in all-atom MD simulations.

#### DNAJB6b side chain virtual bead position

The position of the side chain virtual bead is based on the 3fd virtual site definition in GROMACS^62^. In this 3fd scheme, the position of the virtual site (**r**_vs_) is defined by

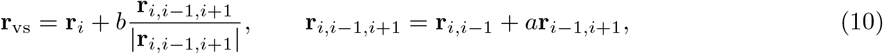

where the virtual bead position is defined using parameters *a* and |*b*|, which control the in-plane angle and distance from the central massive bead (backbone bead), respectively (Eq. 10). A detailed explanation of the coordinate construction is provided in the SI, along with an illustration in Fig. 1, the section *Side chain: 3fd virtual site* in the SI, and Supplementary Fig. 2.

The parameters *a* and |*b*| for virtual side chain placement were calibrated for each residue type in DNAJB6b using the data from all-atom MD simulations^42^ (see supplementary section *All-atom MD simulations*). For this, the center-of-mass positions of side chains and the corresponding C_α_ atom positions were used to compute the geometric parameter *a* and |*b*| for the amino acids in DNAJB6b, revealing that side chain geometry is strongly influenced by the residue’s secondary structure (Fig. 2c, Supplementary Tab. 2, Supplementary Fig. 4–21). To account for this, the data were categorized into four secondary structure types: helix, sheet, coil, and turn and averaged for each structure type (see Supplementary Tab. 2) Each residue was then assigned *a* and |*b*| values based on both its amino acid type and its local secondary structure in DNAJB6b, as determined from all-atom MD simulations (see Supplementary Tab. 1 and 2).

#### DNAJB6b hydrophobicity

For the backbone beads, the hydrophobic scaling constant (α_BB_) was chosen to be 0.22. However, the side chain hydrophobic scaling constant (α_SC_) was calibrated by comparing the radius of gyration (*R*_g_) from CGMD simulations to all-atom MD data for three disordered protein segments: Nsp1 (low charge/hydrophobicity, C/H ratio), Nup60 (high C/H ratio), and Nup145 (low C/H ratio) (see supplementary section *All-atom MD simulations* for details). These simulations were carried out using the side chain parameters for the DNAJB6b coil type residues with α_SC_ values ranging from 0.22 to 0.32 in steps of 0.01, each run for 2 µs, with side chain hydrophobicity values taken from 1BPA-1.1^50^. Based on the variation of *R*_g_ with α_SC_ for Nsp1, Nup145, and Nup60 (Fig. 2b), the optimal value for 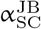 was found to be 0.28, which yielded the lowest error and was thus selected as the hydrophobic scaling factor for DNAJB6b (see Table 1).

**Table 1:**
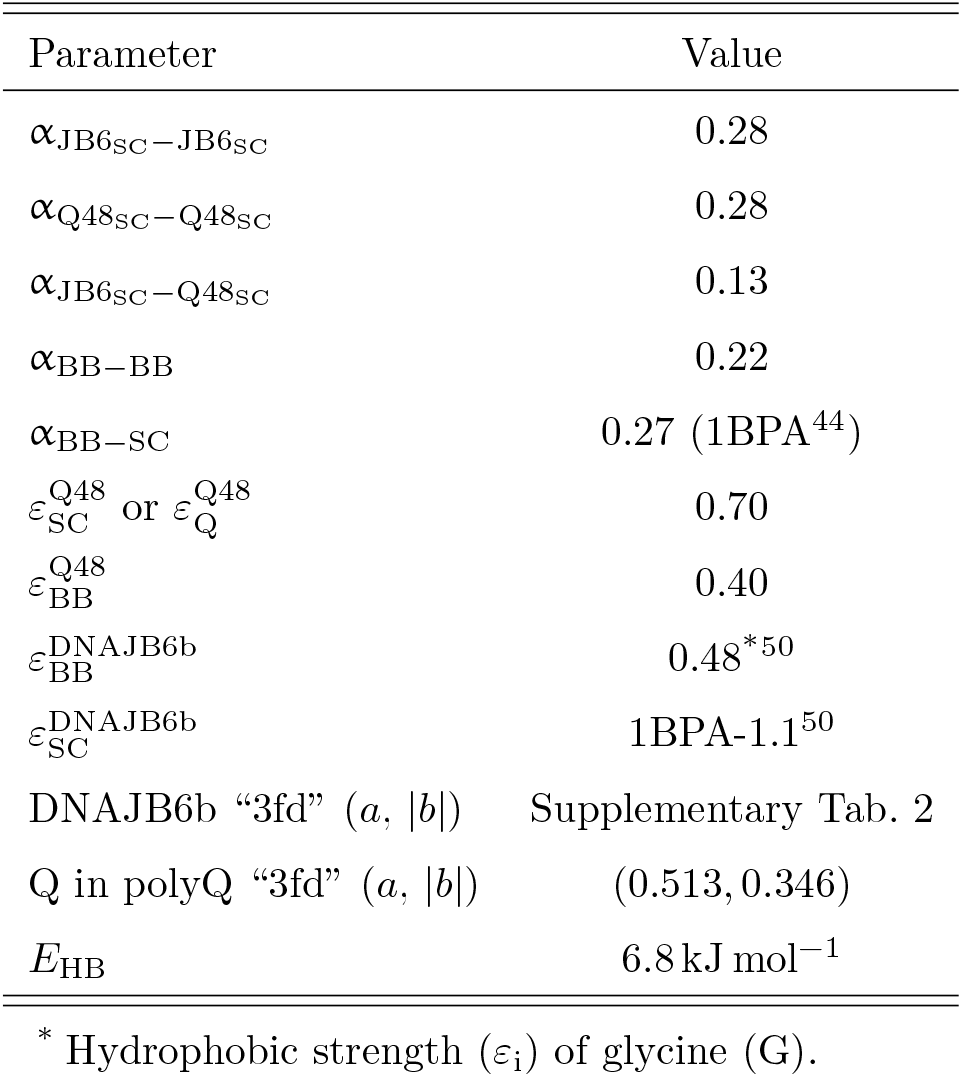
Final parameters for the 2BPA-HB CGMD model. The final parameters used in the DNAJB6b-polyQ and polyQ simulations.

#### Validation of CGMD DNAJB6b against all-atom MD data

With the geometry and hydrophobic parameters of DNAJB6b calibrated, we performed single-molecule CGMD simulations and compared the results to all-atom MD reference data. Satisfyingly, the radius of gyration for the three distinct conformational states of DNAJB6b closely matched the all-atom MD simulations, although the all-atom data was somewhat overpredicted (Fig. 2d, Supplementary Tab. 3). We also compared the intramolecular contact maps between the CGMD and all-atom MD models. The major contact features observed in the all-atom MD simulations were well preserved in the CGMD model (Supplementary Fig. 24).

#### PolyQ side chain virtual bead position

To choose the *a* and |*b*| values for the glutamines in polyQ, we analysed these values for all-atom MD simulations of the sheet regions of DNAJB6b (see Supplementary Tab. 1), for 20 polyQ molecules forming a condensate, and for an amyloid of polyQ (see supplementary section *All-atom MD simulations*). Interestingly, the |*b*| did not have a significant influence across the three cases, but the *a* parameter is higher for DNAJB6b (0.587, see Supplementary Fig. 8, Supplementary Tab. 2) and polyQ in a condensate (0.568, see Supplementary Fig. 22), but lower in case of the polyQ amyloid (0.513, see Fig. 3). CGMD simulations with higher *a* lead to the formation of twisted amyloid fibrils and occasional β-barrels, rather than linear polyQ amyloid like structures as observed for *a* = 0.513. We therefore selected *a* = 0.513 for the CGMD model.

**Figure 3:**
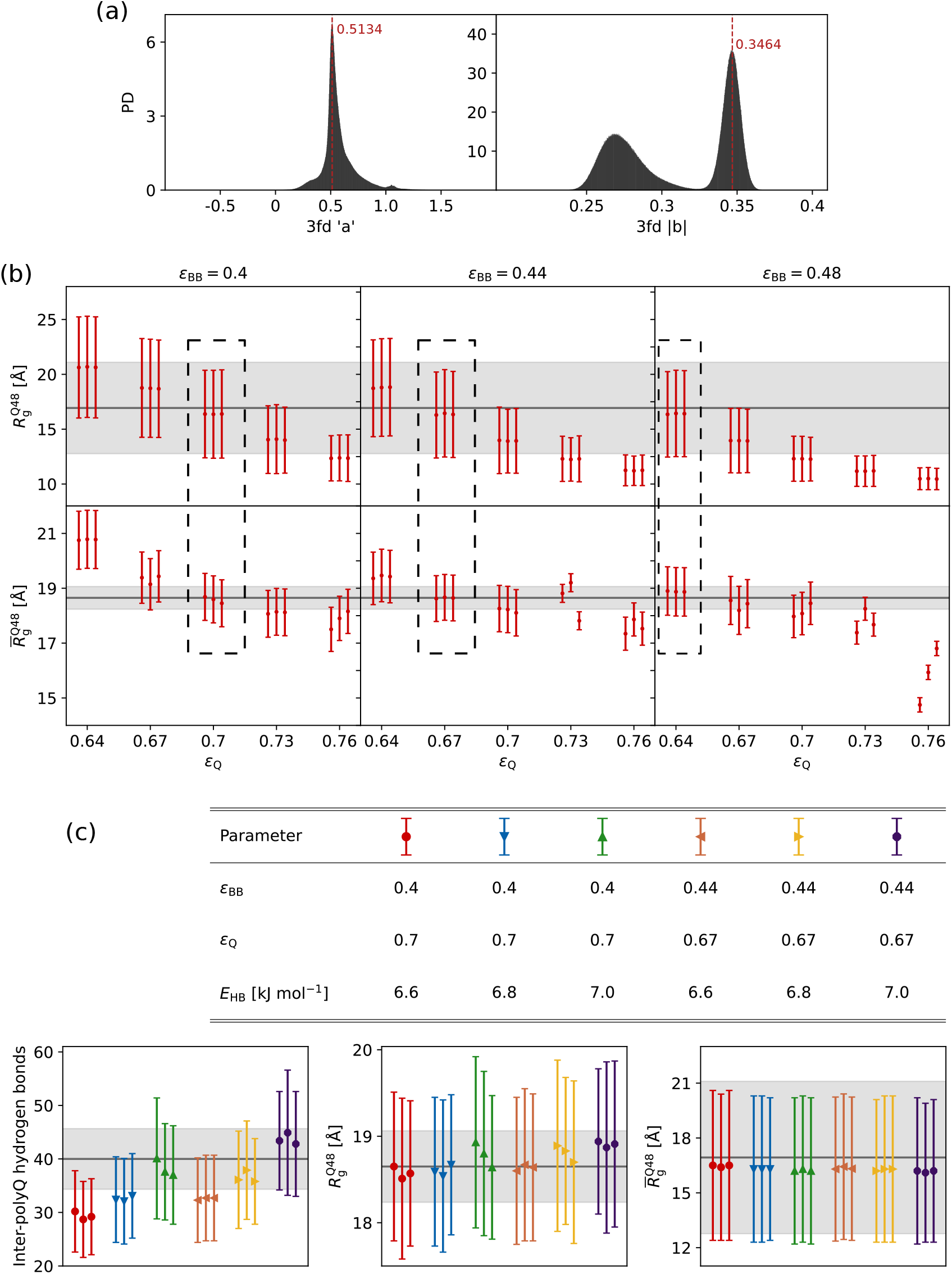
Calibration of parameters for polyglutamine. **(a)** All-atom MD probability density distributions of the 3fd geometry parameters *a* and |*b*| for glutamine, used to calibrate the polyQ. Dashed red lines indicate the dominant mode of each distribution. **(b)** Calibration of glutamine hydrophobicity was performed by comparing the average radius of gyration of a single Q48 molecule 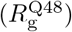 and a 20-molecule Q48 cluster 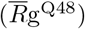 across different values of *ε*_Q_ and *ε*_BB_. **(c)** Summary of calibration results showing inter-polyQ hydrogen bonds in the absence of DNAJB6b, single-molecule *R*_g_, and cluster-averaged *R*_g_ for various combinations of side chain hydrophobicity (*ε*_Q_), backbone hydrophobicity (*ε*_BB_), and hydrogen bonding strength (*E*_HB_). Gray bands in each panel represent all-atom reference values where applicable. Dashed rectangles in **b** show the optimal CGMD parameters where the *R*_g_ matches the all-atom data. Overall, increasing the hydrogen bonding energy results in more inter-polyQ hydrogen bonds (Fig. 3c), while having no significant effect on either the 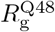 or 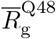. The 3fd parameter *b* has units of length.

#### PolyQ hydrophobicity

To maintain consistency with the DNAJB6b model, the hydrophobicity scaling constants for the back-bone (α_BB-BB_) and side chain (α_SC-SC_) for glutamine were set to 0.22 and 0.28, respectively. For the hydrophobic constants we need the *ε*_*i*_ values (see Eq. 4). For polyQ we only have two of these values: for the backbone (*ε*_BB_) bead and the side chain virtual bead (*ε*_SC_). Calibration of these hydrophobicities was performed by radius of gyration data from all-atom MD simulations (see supplementary section *All-atom MD simulations*) of both a single Q48 molecule 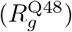 and a 20-molecule Q48 cluster (average 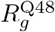, i.e., 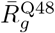, computed over the entire simulation).

CGMD simulations of a single Q48 molecule and a 20-molecule Q48 cluster were performed by varying *ε*_BB_ (0.40, 0.44, 0.48) and *ε*_Q_ from 0.64 to 0.76 in steps of 0.03. 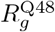 and 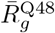 are calculated and compared with all-atom data as shown in Fig. 3b and found three sets to represent the all-atom data well: (1) *ε*_Q_ = 0.7 with *ε*_BB_ = 0.4, (2) *ε*_Q_ = 0.67 with *ε*_BB_ = 0.44, and (3) *ε*_Q_ = 0.64 with *ε*_BB_ = 0.48.

#### PolyQ hydrogen bonding energy

The hydrogen bonding energy (*E*_HB_) is a crucial parameter for the transition of disordered polyQ molecules to an amyloid like structure. Two measures were implemented to determine the optimal hydrogen bonding energy: (1) the number of inter-polyQ hydrogen bonds in the disordered polyQ cluster, and (2) the frequency of amyloid formation. For every set of *ε*_BB_ and *ε*_Q_, the hydrogen bonding energy was varied from 6.6 kJ mol^−1^ to 7.0 kJ mol^−1^ with a step of 0.2 kJ mol^−1^. CGMD simulations were performed using the above parameters to calculate intermolecular polyQ hydrogen bonds, as well as 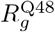 and 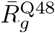. The results show that *R*_g_ remains largely unaffected by changes in *E*_HB_, while increasing *E*_HB_ leads to a higher number of hydrogen bonds between polyQ molecules in a cluster (Fig. 3c). To further optimize the parameters, the frequency of amyloid formation was considered where CGMD simulations were performed with 160 Q48 molecules. Three replica simulations were conducted using two sets of hydrophobicity parameters, with the hydrogen bond energy parameter (*E*_HB_) varied from 6.6 kJ mol^−1^ to 7.0 kJ mol^−1^. The lowest *E*_HB_ at which amyloid formation was observed was for the parameters *E*_HB_ = 6.8 kJ mol^−1^, *ε*_Q_ = 0.70 (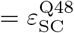 in table 1) and *ε*_BB_ = 0.40 (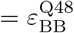 in table 1) (Supplementary Fig. 25), which are chosen as the final parameters for polyQ hydrophobicity and hydrogen bonding. For DNAJB6b the same parameters as polyQ were used for hydrogen bonding.

#### Calibration of interactions between DNAJB6b and polyQ

The final parameter calibrated was the hydrophobicity scaling factor (α_JB6-Q_), which governs the interaction strength between DNAJB6b and polyQ. All-atom MD simulations of five DNAJB6b molecules with 20 Q48 peptides, starting from the solution, were used as reference for this calibration (see supplementary section *all-atom simulations*), where multiple molecules of DNAJB6b were added to a solution of polyQ peptides. In these all-atom MD simulations we observed that the DNAJB6b and polyQ molecules co-condensate with the chaperone directly binding to polyQ and reducing the polyQ-polyQ hydrogen bonding required for amyloid formation.

#### Validation of CGMD DNAJB6b-polyQ interactions against all-atom data

Using the CGMD parameters obtained (detailed in the sections above), CGMD simulations were conducted for systems containing 20 Q48 and five DNAJB6b molecules, initialized either from solution or a preformed polyQ cluster. Contact analysis revealed that the S/T-rich and G/F_1_-rich regions are the primary regions interacting with polyQ, consistent with all-atom MD results (Fig. 4c, Supplementary Fig. 27). Notably, the interaction profiles of DNAJB6b with polyQ molecules remained similar across simulations starting either from the solution or the disordered aggregate, a feature not captured in all-atom MD simulations due to limited sampling (Supplementary Fig. 27). Amino acid–level contact analysis further confirmed strong agreement between the CGMD and all-atom MD models (Fig. 4d, Supplementary Fig. 28).

**Figure 4:**
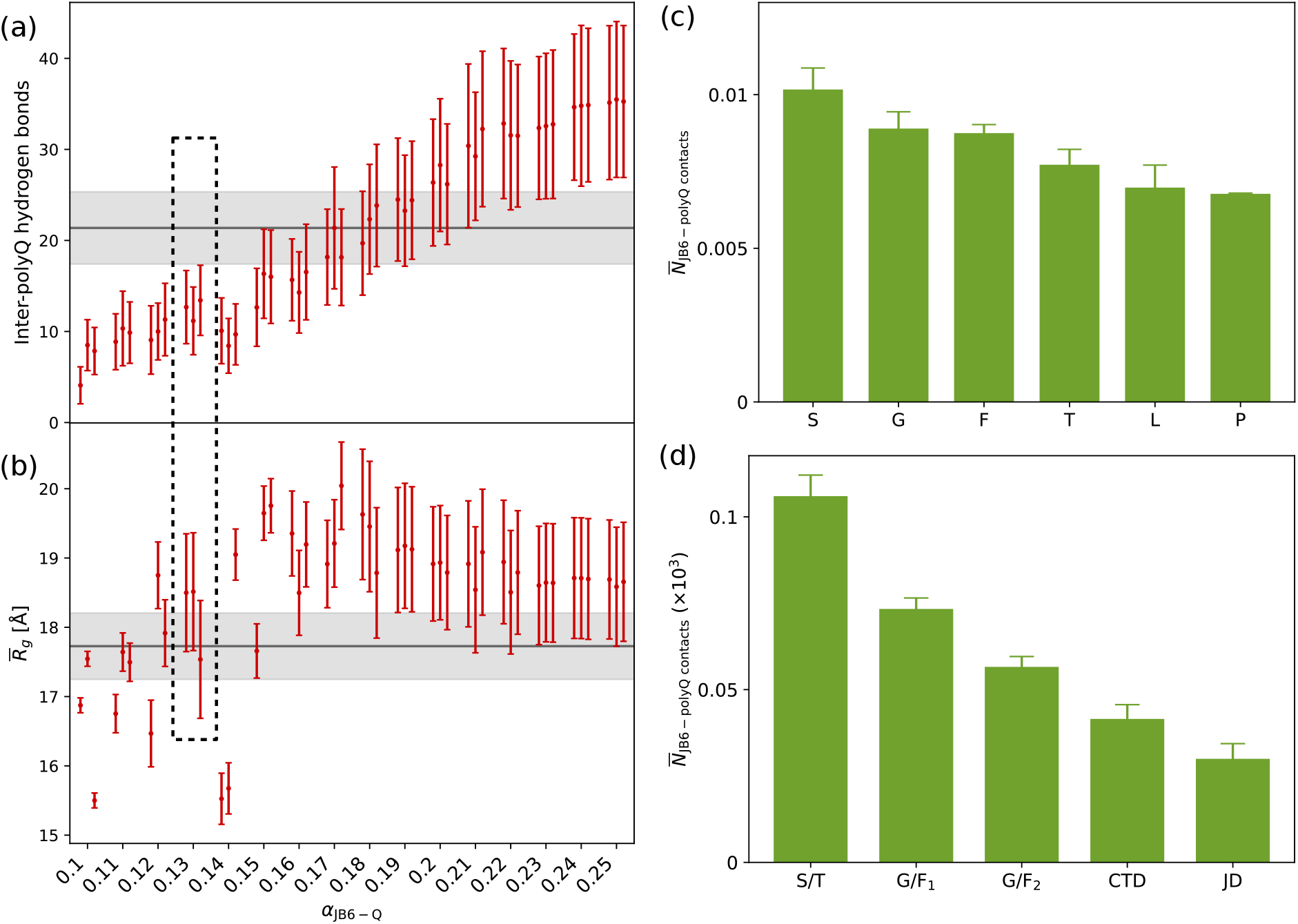
Calibration of parameters for the interaction strength between polyglutamine and DNAJB6b. **(a)** and **(b)** Variation of the number of inter-polyQ hydrogen bonds and the average radius of gyration 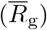 of 20 Q48 molecules in solution in the presence of five DNAJB6b molecules, across different values of the interaction scaling factor α_JB6-Q_. The dashed rectangle indicates the optimal parameter region, where α_JB6-Q_ = 0.13 was selected. Gray bands in each panel (a) and (b) represent the corresponding all-atom MD reference values. **(c)** and **(d)** Domain-level and amino acid–level interaction profiles of DNAJB6b with polyQ, respectively, obtained from CGMD simulations of 20 Q48 and five DNAJB6b molecules using the final calibrated parameters (Tab. 1). These results are consistent with all-atom MD simulations; detailed comparison is shown in Supplementary Fig. 27, 28.

To capture this behaviour observed in all-atom MD, CGMD simulations were performed for 20 Q48 peptides and five DNAJB6b starting from solution with α_JB6-Q_ values ranging from 0.11 to 0.25 in steps of 0.01. For α_JB6-Q_ *>* 0.20, no significant reduction was observed in the number of inter-polyQ hydrogen bonds and the average radius of gyration 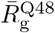 compared to simulations without DNAJB6b (Fig. 4a, 4b). Since all-atom data showed that DNAJB6b and polyQ co-condense and reduce the number of intermolecular hydrogen bonds between polyQ peptides (*N*_HB_), CGMD simulations with an α_JB6-Q_ of 0.13 reproduced both the correct 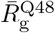 and a lower number of inter-polyQ hydrogen bonds, and was therefore used in subsequent simulations (Fig. 4a, 4b). The final CGMD parameters are provided in table 1.

#### Final parameters

The final parameter set for the 2BPA-HB coarse-grained model used in the DNAJB6b–DNAJB6b, DNAJB6b–polyQ and polyQ-polyQ simulations comprises calibrated hydrophobic scaling factors, residue-type-dependent hydrophobic strengths, virtual-site geometry parameters, and a hydrogen-bond energy tuned to reproduce amyloid formation. For hydrophobic scaling, the backbone scaling was set to α_BB−BB_ = 0.22, and the backbone–side-chain scaling was fixed to α_BB−SC_ = 0.27 as in the 1BPA model. Side-chain scaling within DNAJB6b and within Q48 was set to 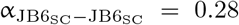 and 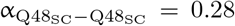, while the cross interaction between DNAJB6b and Q48 side chains was tuned to 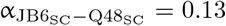 to reproduce co-condensation and the reduction of inter-polyQ hydrogen bonding observed in all-atom simulations. PolyQ hydrophobic strengths were chosen as 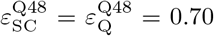 and 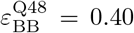, whereas DNAJB6b uses 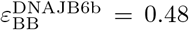, and 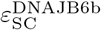 taken from the 1BPA-1.1 parameter set. Side-chain placement follows the GROMACS “3fd” scheme, with DNAJB6b residue- and secondary-structure-dependent (*a*, |*b*|) values listed in Supplementary Tab. 2, and glutamine in polyQ fixed to (*a*, |*b*|) = (0.513, 0.346). Hydrogen bonding was modeled using the shifted Lennard–Jones form with *E*_HB_ = 6.8 kJ mol^−1^, applied consistently for both DNAJB6b and polyQ. A simplified flowchart describing the calibration process of the 2BPA-HB model is provided in the supplementary text, Supplementary Fig. 23.

## Results

### Effect of DNAJB6b on the polyglutamine aggregation pathway

Simulations using the CGMD model show that polyQ molecules initially associate into structurally disordered aggregates and can subsequently reorganize into ordered, fibril-like amyloid structures (Supplementary Fig. 25)^56^. This aggregation pathway provides three stages at which DNAJB6b may exert its anti-aggregation activity: (I) when the polyQ molecules are dispersed in solution, (II) after a disordered aggregate has formed, and (III) after an ordered amyloid-like structure has formed. To evaluate the chaperone’s effectiveness across these stages, we performed simulations with a system containing 13 DNAJB6b molecules (5 closed, 5 open and 3 extended conformations) and 52 polyQ molecules at various stages.

Control simulations without DNAJB6b show that the number of intermolecular polyQ hydrogen bonds (*N*_HB_) stabilizes at an average of 91 ± 4 (Fig. 5a). Introducing DNAJB6b into a solution of polyQ molecules results in a substantial reduction in *N*_HB_, with an average value of 21 ± 1, corresponding to a 78 % decrease (Fig. 5a, 5c). When simulations are initiated from a preformed polyQ disordered aggregate, *N*_HB_ decreases to 59 ± 10, which is higher than in the polyQ solution case, but still significantly lower than the control (Fig. 5a). Interestingly, DNAJB6b molecules are able to penetrate the disordered aggregate of polyQ, with the serine/threonine-rich (S/T) region showing the greatest degree of insertion (Fig. 5c, 5d, and Supplementary Fig. 26) which contrasts with our earlier all-atom MD simulations (Adupa *et al. In preparation*), where such penetration was not observed due to sampling limitations in the all-atom MD. In simulations starting from a preformed polyQ amyloid fibril (see supplementary section *all-atom simulations*), the average *N*_HB_ in the absence of chaperone is 858 ± 17 (Fig. 5b). Upon addition of DNAJB6b, this value decreases to 644 ± 13 (Fig. 5b, 5h). In this case, DNAJB6b forms a coating around the fibril surface (Fig. 5h), and the observed reduction in *N*_HB_ is primarily due to partial disruption of less stable regions of the amyloid, particularly where only a few polyQ strands interact. This structural change can be visually appreciated by comparing the amyloid before and after chaperone addition (Fig. 5h and 5i).

**Figure 5:**
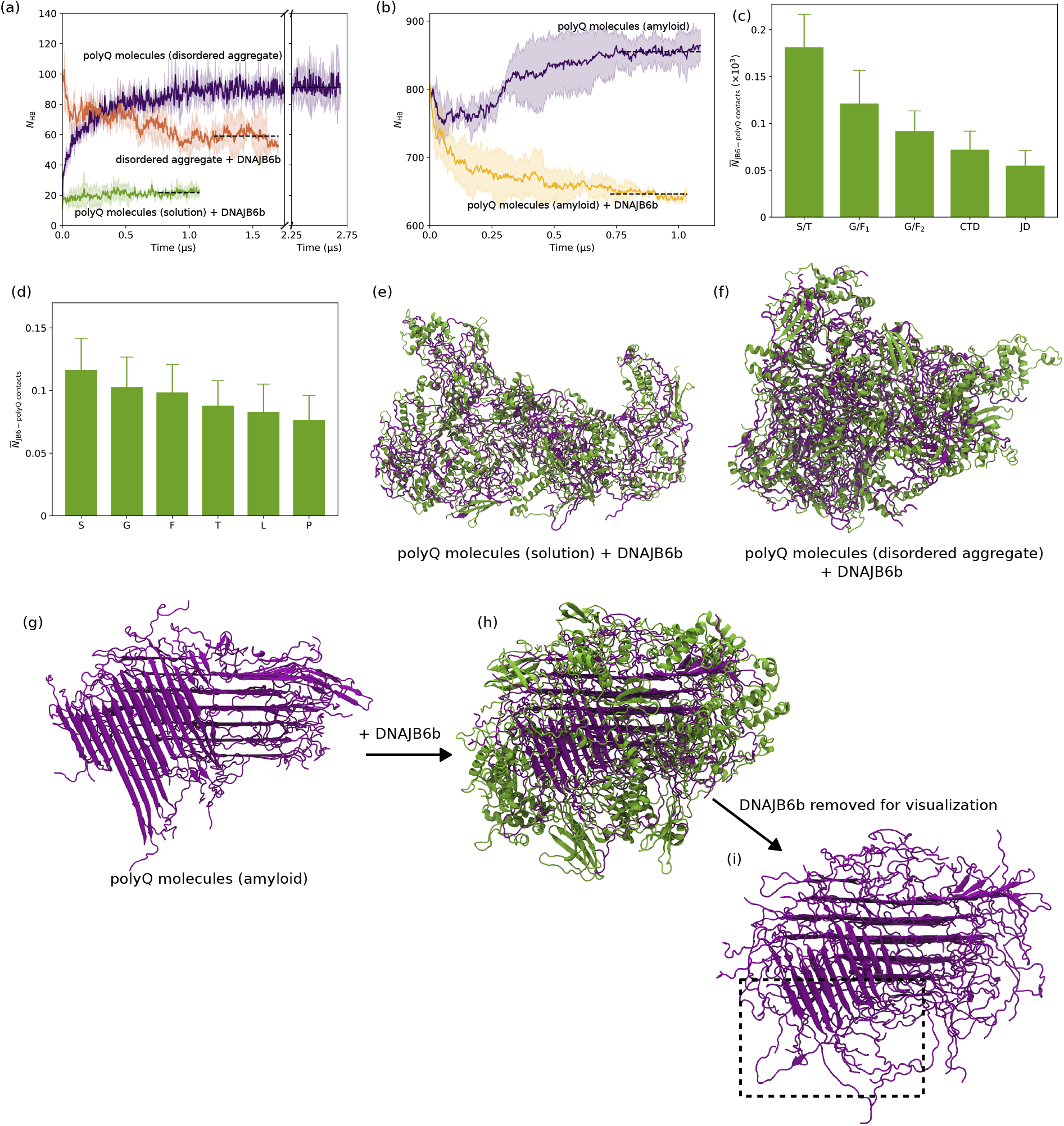
Effect of DNAJB6b on polyglutamine aggregation at various configurations of Q48 peptides. Plots showing the variation in the number of intermolecular polyQ hydrogen bonds over time (*N*_HB_) for a system of 52 Q48 peptides with 13 DNAJB6b molecules. **(a)** Simulations where polyQ molecules are in solution without DNAJB6b (purple), in the presence of DNAJB6b (green), and DNAJB6b added to a pre-existing disordered aggregate of polyQ (orange). **(b)** Simulations where polyQ molecules form an amyloid from a solution of polyQ molecules in the absence of DNAJB6b (purple), and after the addition of DNAJB6b at t=0 (yellow). **(c), (d)** Domain and amino acid level interaction profiles of DNAJB6b with polyQ peptides, respectively, obtained from CGMD simulations of 52 Q48 and 13 DNAJB6b molecules starting from a solution using the final calibrated parameters (Tab. 1). Snapshots show the 52 polyQ peptides + 13 DNAJB6b molecules in their final configurations for **(e)** polyQ in solution with DNAJB6b added, and **(f)** DNAJB6b added to a preformed disordered aggregate of polyQ. The structure of the polyQ amyloid is shown **(g)** before addition of DNAJB6b, **(h)** after addition where DNAJB6b coat the amyloid surface, and **(i)** highlighting the effect of DNAJB6b on the fibril. For the plots in **(a), (b)**, data from all three replicas were included, and a running average was applied using a window size of 2.5 ns.

### Impact of DNAJB6b conformations on the polyQ aggregation stages

Our previous all-atom MD study demonstrated that DNAJB6b is a dynamic molecule that transitions between three conformational states: closed, open, and extended^42^. In this work, we leveraged the ability of the CGMD model to sample these conformational transitions and performed simulations with DNAJB6b initialized in specific states to examine their effect on polyglutamine aggregation.

While the previous section focused on simulations with a mixed ensemble of DNAJB6b (closed:open:extended = 5:5:3), here we present results from simulations where the chaperone was initialized exclusively in one of the three conformations: closed (13:0:0), open (0:13:0), or extended (0:0:13). We tested the effect of DNAJB6b conformations on polyQ aggregation at three distinct stages: solution, disordered aggregate, and amyloid fibril (Fig. 6a-c). While only minor differences in inter-polyQ hydrogen bonding (*N*_HB_) were observed across stages of polyQ aggregation, the disordered aggregate showed the most variation. Here, the effectiveness of DNAJB6b appeared to depend on the degree of exposure of the S/T-rich region, which is partially accessible in the open state and fully exposed in the extended state. Notably, the open conformation can transition into the extended state during simulation, further increasing S/T-rich region accessibility (Supplementary Tab. 4). In this configuration, the S/T-rich region penetrates into the disordered aggregate of polyQ, while the remainder of the DNAJB6b molecule remains at the surface (Fig. 5c, Supplementary Fig. 26). Overall, these findings suggest that DNAJB6b’s anti-aggregation activity is largely robust across conformational states, with subtle state-dependent differences emerging primarily in the disordered aggregate context.

**Figure 6:**
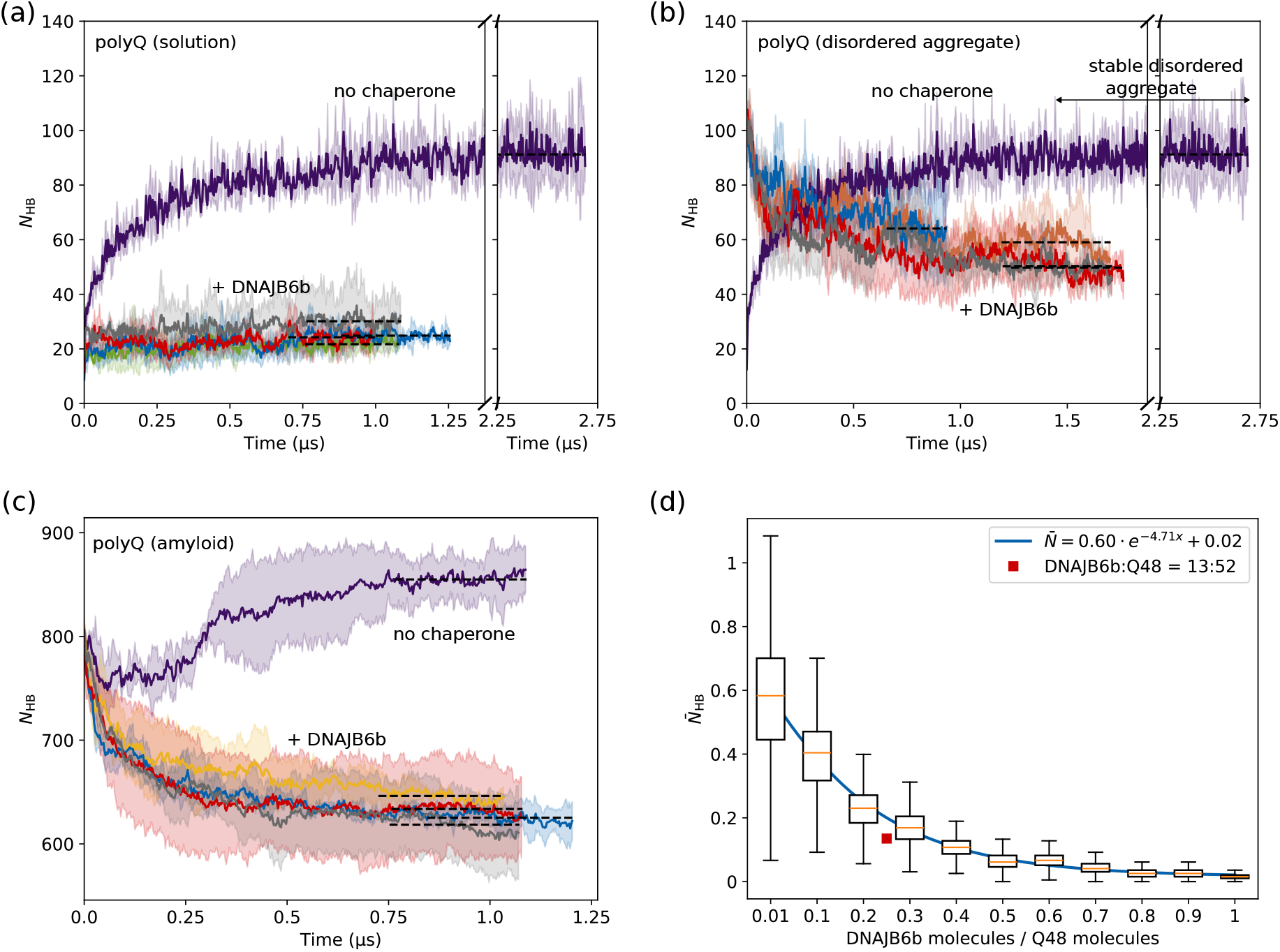
Effect of conformational states of DNAJB6b on polyglutamine aggregation at various configurations of Q48 peptides. Time evolution of the number of intermolecular hydrogen bonds between polyQ molecules (*N*_HB_) under different conditions. **(a)** Simulations starting from a polyQ solution, without DNAJB6b or with DNAJB6b added in different conformational ensembles: a mixed set (five closed, five open, and three extended), exclusively closed, exclusively open, or exclusively extended. Simulations initialized from a preformed polyQ cluster, followed by the addition of DNAJB6b in the same conformational ensembles. **(c)** Simulations initialized from a preformed polyQ amyloid, with DNAJB6b added in the same conformational ensembles. For all cases, 13 DNAJB6b molecules are added, where color code represent: mixed DNAJB6b (green, orange and yellow in panels **(a), (b)** and respectively), exclusively closed state (blue), exclusively open state (red), exclusively extended state (gray). PolyQ only simulations are shown in purple. For the plots in **(a), (b)** and **(c)**, data from all three replicas were included, and a running average was applied using a window size of 2.5 ns. **(d)** Box plot showing the normalized number of intermolecular hydrogen bonds (*N*_HB_) between polyQ molecules as a function of the DNAJB6b-to-Q48 ratio. Normalization was performed using the average *N*_HB_ from simulations where polyQ molecules in solution spontaneously formed a cluster (195.5). An exponential decay fit is shown in blue, and the average result from simulations with 52 Q48 molecules and 13 openstate DNAJB6b molecules is indicated by the red square. The system consists of 100 polyQ (Q48) molecules with varying numbers of DNAJB6b molecules: 1, 10, and then increasing in steps of 10 up to 100.

### Effect of DNAJB6b mutations on polyglutamine aggregation

The contact analyses identified the S/T-rich and G/F-rich regions as the main regions of DNAJB6b interacting with polyQ (Fig. 4d, 5c). To examine whether the sequence composition of these regions contributes to the suppression of polyQ aggregation, we compared wild-type DNAJB6b with the M3 (18 S/T→A), the 12 F→A mutant in the S/T domain, and the G/F→A mutant in the G/F_1_ domain.

All three mutants showed an increase in the number of inter-polyQ hydrogen bonds compared with wild-type DNAJB6b (Supplementary Fig. 29). The median number of inter-polyQ hydrogen bonds increased from approximately 22 for wild-type DNAJB6b to 31 for M3 and to 37 for both the 12 F→A and G/F→A mutants. The reduced activity of the S/T-rich and G/F-rich mutants supports a sequence- and domain-dependent contribution of these regions to the suppression of polyQ aggregation in line with the experimental work from Kakkar *et al*.^30^.

### Chaperone efficiency at varying DNAJB6b:polyQ ratios

Experimental studies have shown that DNAJB6b remains effective even at sub-stoichiometric ratios relative to its client proteins. To investigate this, we performed CGMD simulations with 100 Q48 molecules and varying numbers of DNAJB6b molecules (1, 10, 20 … 100), using exclusively the open conformation. All simulations were initiated from polyQ in solution, allowing us to assess the chaperone’s effectiveness across a wide range of DNAJB6b:Q48 ratios.

For each DNAJB6b:Q48 ratio, the number of intermolecular hydrogen bonds between polyQ molecules was calculated and normalized using the average *N*_HB_ from a polyQ-only disordered aggregate simulation (195.5). The resulting normalized values are shown as a box plot in Fig. 6d, which reveals that beyond a DNAJB6b:Q48 ratio of 0.5, further reductions in inter-polyQ hydrogen bonding plateau, suggesting that each DNAJB6b molecule can effectively handle up to two polyQ molecules before reaching saturation. To quantify the relationship between chaperone concentration and polyQ aggregation, an exponential decay function was fitted to the data.

## Discussion

In this study, we developed a two-beads-per-amino-acid (2BPA-HB) CGMD model to investigate polyglutamine aggregation and the role of the molecular chaperone DNAJB6b in modulating this process. The model was primarily parameterized using data from all-atom molecular dynamics simulations. Using the 2BPA-HB model, we observed that Q48 molecules initially associate into structurally disordered aggregates and subsequently reorganize into ordered amyloid-like structures. This behaviour supports a two-stage disorder-to-order aggregation pathway under the simulated conditions, where polyQ molecules initially undergo oligomerization into a disordered aggreagate, followed by a disordered-to-order phase transition in which the disordered aggregate converts into an ordered amyloid fibril—consistent with observations from experimental studies on synthetic polyQ peptides^65,66^.

Amyloid fibrils are considered the toxic species driving Huntington’s disease, and previous studies have identified molecular chaperones—particularly DNAJB6b—as key suppressors of polyglutamine aggregation^27–31^. However, the molecular mechanism by which DNAJB6b prevents aggregation remains poorly understood.

The primary objective of this study was therefore to mechanistically investigate how DNAJB6b suppresses polyglutamine aggregation. To this end, we calibrated our CGMD model using all-atom simulation data capturing interactions between DNAJB6b and polyQ peptides. Simulations in which DNAJB6b was introduced into a solution of polyQ molecules revealed that the chaperone co-condensates with the Q48 peptides, effectively shielding them from forming intermolecular hydrogen bonds between polyQ molecules, which are essential for the transition to amyloid-like structures. While this suggests that co-condensation plays a role in suppression, the precise mechanism—particularly at the disordered aggregate stage—remains complex.

Previous studies on FUS proteins have shown that amyloid formation within disordered droplets initiates at the droplet surface, where the protein dynamics is the highest^67,68^. In our simulations of Q48 with DNAJB6b, we consistently observed a reduction in the dynamics of polyQ molecules upon interaction with the chaperone. Even in the system containing a single DNAJB6b molecule and a disordered aggregate of 52 Q48 peptides, where no significant reduction in *N*_HB_ was observed, the average root-mean-square fluctuation (RMSF) of the Q48 molecules was reduced (Supplementary Fig. 30–32). Based on this, we hypothesize that at sub-stoichiometric ratios, DNAJB6b suppresses polyQ aggregation by lowering molecular mobility, thereby reducing the likelihood of transitioning into amyloid fibrils.

Our previous all-atom MD study showed that DNAJB6b exists as a dynamic ensemble, transitioning between three conformations: closed, open, and extended. The CGMD 2BPA-HB model developed in this work reproduces these transitions and enables us to probe their relevance to chaperone function. To evaluate the impact of DNAJB6b conformation on polyglutamine aggregation, we performed simulations using the closed, open, and extended states of the chaperone across three polyQ aggregation stages: solution, disordered aggregate, and amyloid. In simulations starting from polyQ in solution or amyloid fibrils, we observed no substantial differences in DNAJB6b’s effectiveness across conformational states. However, when polyQ aggregation was initiated from a disordered aggregate, the open and extended states of DNAJB6b were consistently more effective in suppressing aggregation than the closed or mixed states. This difference appears to be linked to the accessibility of the S/T-rich region, which is partially exposed in the open state and fully exposed in the extended state. The high exposure of the S/T-rich region in the open and extended states facilitates its insertion into the disordered aggregate of polyQ, unlike in the closed state where the domain remains collapsed. This limitation is less relevant when the polyQ molecules start in solution, where the dynamic fluctuations of the polyQ molecules are sufficiently large to create multivalent interactions between the S/T region and polyQ residues. This is not the case for the disordered aggregates where the open and extended dynamics enhances penetration compared to the closed state.

The mutation simulations further show that DNAJB6b activity cannot be explained by nonspecific association with polyQ alone. Removing the residue-specific interactions of the M3 S/T mutant, the phenylalanine residues in the S/T-rich region, or the glycine and phenylalanine residues in the G/F_1_-rich region increased inter-polyQ hydrogen bonding. Together with the contact profiles, these results indicate that both the S/T-rich and G/F-rich regions contribute to the interaction with polyQ and to the suppression of polyQ aggregation.

The relatively simple representation of 2BPA-HB provides computational efficiency but also limits the scope of the model. The use of a single massive backbone bead, together with virtual side-chain and hydrogen-bonding sites, is sufficient for the aggregation behaviour and DNAJB6b–polyQ interactions examined here. However, an additional backbone bead may become necessary when a more detailed description of peptide-plane geometry, local secondary-structure formation, competing hydrogen-bonding arrangements, or amyloid polymorphism is required. In addition, the present model employs a fixed side-chain geometry and therefore does not capture alternative side-chain conformations, such as the type A and type B glutamine conformations reported for polyQ amyloids^69^. Models such as AWSEM-IDP follow a complementary parameterization philosophy, making greater use of statistical and energy-landscape-based information^52,53^. Future developments could combine atomistically informed local chemical specificity with such statistical approaches to improve transferability and global conformational and aggregation behaviour.

In conclusion, our study presents a CGMD model that captures key features of polyglutamine aggregation and its modulation by the molecular chaperone DNAJB6b. The model reproduces essential aspects of the polyglutamine aggregation process, including phase separation and amyloid fibril formation, while allowing access to system sizes and timescales beyond the reach of all-atom simulations. Using this framework, we demonstrate that DNAJB6b inhibits polyglutamine aggregation through co-condensation and thereby reduces the dynamics of polyQ molecules that together help explain its effectiveness, even at sub-stoichiometric ratios. By further enabling the exploration of conformational states of DNAJB6b, the developed 2BPA-HB CGMD model offers a powerful foundation for future studies aimed at uncovering the molecular principles of chaperone-mediated aggregation suppression and potential therapeutic strategies for Huntington’s disease.

## Notes

### Competing Interest Statement

The authors have declared no competing interest.

### Funding

- Nederlandse Organisatie voor Wetenschappelijk Onderzoek

