## Supplementary Information for "A two-bead-per-aminoacid coarse-grained MD model with hydrogen bonding (2BPA-HB) to probe DNAJB6b-mediated suppression of polyglutamine aggregation in Huntington’s disease"

#### All-atom simulations

The interaction strength ($\alpha$) between the SC-SC beads of either DNAJB6b or Q48 (see Fig. 1) are calibrated based on FG-Nup segments (Nsp1, Nup60, and Nup145) all-atom explicit water MD simulations, which were performed using the GROMACS MD package, version 2021.3[1]. Simulations were performed using the CHARMM36m forcefield with TIP3P water[2]. A leap-frog integrator with a timestep of 2 fs was used to integrate Newton's equations of motion. Short-range electrostatic and van der Waals interactions were truncated at 1.0 nm, while long-range electrostatics were computed using the particle mesh Ewald (PME) method with a grid spacing of 0.125 nm and a PME order of four. Temperature was maintained at 300 K using a velocity-rescaling thermostat with a time constant of 0.1 ps[3], and pressure was controlled at 1 bar using the Parrinello–Rahman barostat[4] and an isothermal compressibility of $4.5 \times 10^{-5}$ bar$^{-1}$. All bonds involving hydrogen atoms were constrained using the LINCS algorithm[5]. Initially, each FG-Nup segment was placed at the center of a cubic box with a 1.0 nm padding and solvated with water. $Na^+$ and $Cl^-$ ions were added by replacing water molecules to achieve a salt concentration of 100 mM. The systems were first energy minimized using the steepest descent algorithm. This was followed by two consecutive equilibration runs: a 1 ns NVT ensemble simulation to stabilize temperature, and a 1 ns NPT simulation to equilibrate density. Finally, the production simulations were then carried out in the NPT ensemble for at least 1 µs.

The interaction strength ($\alpha$) between the SC-SC beads of DNAJB6b and Q48 (see Fig. 1) are calibrated based on all-atom explicit water MD simulations of single Q48 and 20 Q48 peptides. The methodology descibed above is also used for conducting these simulations.

The virtual bead position for side chains in DNAJB6b, i.e. the `3fd` ($a$, $|b|$) and `3out` virtual site parameters, were calibrated using previously published single-molecule all-atom simulations of DNAJB6b[6].

The calibration of polyQ side chain virtual bead position was carried out in two steps: (1) an amyloid fibril was first generated through a CGMD simulation of 52 Q48 molecules starting from solution, using a higher hydrogen-bonding energy ($E_{\mathrm{HB}}$) of 7.2 kJ mol$^{-1}$; (2) the resulting fibril was backmapped to an all-atom structure, overlaps were removed, and an all-atom simulation was performed for 500 ns. The final structure obtained was then used to calibrate the `3fd` ($a$, $|b|$) parameters of polyQ.

#### Coarse-Grained simulations

Following this, we conducted 2BPA simulations of single-molecule DNAJB6b, single-molecule polyQ (Q48) peptides, multiple-molecule Q48 peptides, and multiple-molecule DNAJB6b-Q48 using the 2BPA model, systematically testing various combinations of BB and SC interaction strengths to identify the combination that best captures the single-molecule behaviour of the molecular chaperone DNAJB6b, clustering behaviour of polyQ peptides, and the anti-aggregation behaviour of DNAJB6b on Q48 aggregation. CGMD simulations were performed with the GROMACS[1] software package (version 2019.6), with a modified implementation of the `3out` virtual site scheme[7] to construct the HB beads as defined in Eq. 3. Simulations are conducted at 300 K using a time step of 15 fs and inverse friction coefficient of $\gamma^{-1}$ of 2 ps for the Langevin dynamics integrator.

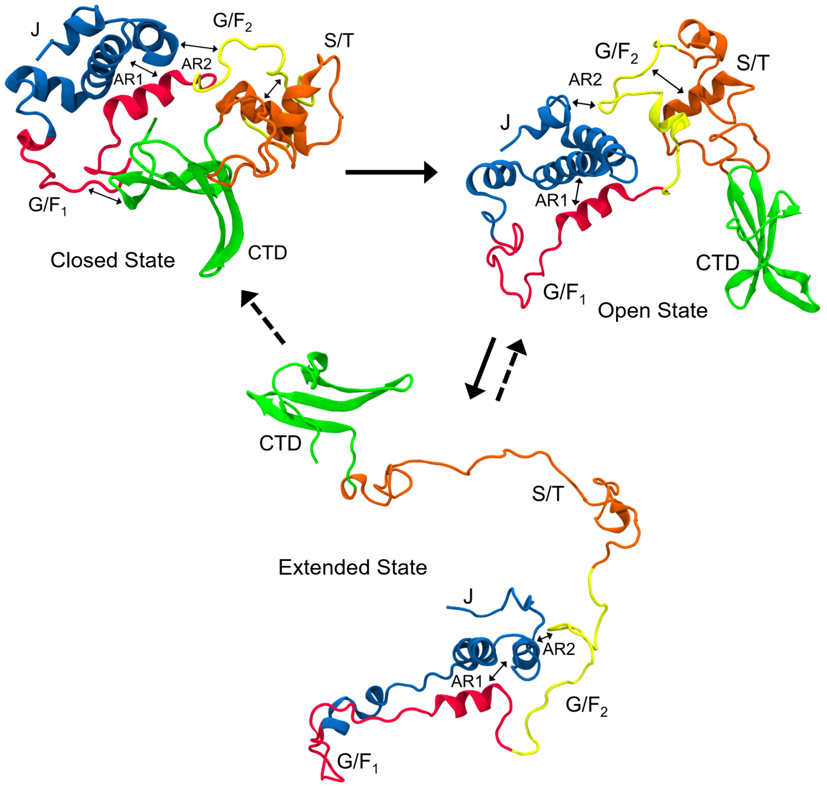


**Supplementary Figure 1: Conformational states of DNAJB6b.** The co-chaperone DNAJB6b in its three tertiary structure states. The possible transitions between the states are shown with arrows. The double-sided arrows show the intra-molecular interactions between domains. Figure adapted from Adupa *et al.*[6]

#### DNAJB6b mutant simulations

To examine the contribution of the S/T-rich and G/F-rich regions to DNAJB6b activity, we performed simulations with three DNAJB6b mutants. The M3 mutant contains substitutions of 12 S and six T residues in the S/T-rich region, the 12 F→A mutant contains substitutions of all 12 F residues in the S/T-rich region, and the G/F→A mutant contains substitutions of all nine G and eight F residues in the $G/F_1$-rich region. In the CG model, the substituted positions retained their excluded-volume interactions and elastic-network restraints, while their hydrophobic, electrostatic, and hydrogen-bonding interactions were removed. These simulations therefore represent the effect of removing the residue-specific interactions at the mutated positions rather than an exact coarse-grained representation of alanine to maximize the effect of the mutation on the function of DNAJB6b.

Each system contained 13 open-state DNAJB6b molecules and 52 Q48 molecules initially dispersed in solution. Three replica simulations of at least 1 µs were performed for each mutant and for wild-type DNAJB6b. The number of intermolecular hydrogen bonds between Q48 molecules was calculated over the final quarter of each trajectory.

#### Elastic network

##### Secondary structure bonds

To preserve the secondary structure in the 2BPA model of DNAJB6b, an elastic network was applied based on all-atom MD simulations of a single-molecule DNAJB6b[6]. An elastic network consists of a series of extra harmonic bonds that are introduced to restrain the distance between two backbone beads. The harmonic potential is defined by

$$U_{\mathrm{H}}^{ij} = \frac{1}{2} k_{ij} (r_{ij} - b_{ij})^2, \tag{1}$$

Here, $k_{ij}$ is the force constant in kJ mol$^{-1}$, defining the stiffness of the elastic bond, while $r_{ij}$ and $b_{ij}$ denote the instantaneous and equilibrium distances between particles $i$ and $j$, respectively. In the elastic network, a uniform force constant of 8000 kJ mol$^{-1}$ is used.

Using the all-atom trajectory, secondary structure assignments were performed using STRIDE, which classifies residues into seven categories: coil, turn, 3–10-helix, π-helix, α-helix, β-bridge or β-sheet. Residues with over 50% coil or turn content were considered disordered and excluded from the elastic network. Elastic restraints were applied only to stretches of three or more consecutive ordered residues. Supplementary table 1 lists the residues included in the secondary structure elastic network for the closed state.

To maintain the secondary structure of the DNAJB6b, elastic restraints were implemented in the 2BPA model. All-atom MD simulations of single-molecule DNAJB6b[6] were used to determine the secondary structure sections of DNAJB6b. To do this, STRIDE was used for on the DNAJB6b trajectory, where each residue was assigned one of the seven different secondary structures: coil, turn, 3–10-helix, π-helix, α-helix, β-bridge or β-sheet. Coil and turn residues are considered as disordered residues. To determine which residues to include in the elastic network, the percentage of coil and turn structure is used. If the sum of these two is larger than 50%, that residue is considered as disordered, and it will therefore not be included in the secondary structure elastic network. Furthermore, elastic bonds are only added if three or more consecutive residues have less than 50% coil plus turn content. Supplementary Table 1 shows which residues were included in the secondary structure elastic network for the closed state of DNAJB6b.

The elastic-network restraints implemented in this model are used to preserve the ordered structural elements of DNAJB6b while allowing transitions between the open and extended states. However, the closed state was not recovered without an additional restraint, indicating that the model captures only part of the conformational landscape. This limitation should be considered when interpreting the relative stability and interconversion of the three DNAJB6b states.

##### Tertiary structure bonds

While the secondary structure was stabilized, the restraints described previously were insufficient to maintain the full tertiary structure of DNAJB6b during simulations. To address this, a minimal set of additional harmonic restraints (see Supplementary Eq. 1, with stiffness 8000 kJ mol$^{-1}$) was introduced to better capture the dynamics observed in AAMD simulations, while preserving conformational flexibility.

To preserve the tertiary structure of the CTD, additional harmonic restraints were applied between the β-strands. Seven such restraints were introduced between the following residue pairs: K189–V209, V209–R215, R215–T228, T228–E233, T195–T204, T204–E219, and E219–L224. These cross-strand restraints were necessary to prevent strand separation and ensure stability of the β-sheet during simulations.

To maintain the relative positioning of the α-helices in the J-domain, six additional harmonic restraints were applied based on the AAMD simulations of DNAJB6b. These restraints were placed between the following residue pairs: Y4–Y65, Y65–V55, V55–Y4, P16–L56, I19–L56, and V10–A22, connecting helices H1–H4, H3–H4, H1–H3, H2–H3 (twice), and H1–H2 respectively.

One key observation from the all-atom simulations was the stability of the autoinhibitory anchor region (anchor 1) across different conformational states of DNAJB6b. To preserve this feature in the CG model, two elastic restraints were applied between residues A50–P96 and H31–F104. Similarly, to maintain the anchor region 2 (AR2) observed in the all-atom MD simulations, two more restraints were added between residues V10–F123 and A22–F123 were applied in the open and extended states, but omitted in the closed state of DNAJB6b.

With the final parameters (see Table 1), coarse-grained simulations of single DNAJB6b molecules captured the transitions from the closed to open state and from the open to extended state. However, reverse transitions from the open or extended states back to the closed state were rarely observed. To specifically evaluate the behavior of the closed state on polyglutamine aggregation, an additional harmonic restraint was applied between residues E88 and L236—linking the G/F$_1$ domain and the CTD—to prevent transitions to the open or extended conformations. This restraint was omitted in simulations starting from the open or extended states, allowing those molecules to transition freely between conformations.

### Side chain: `3fd` virtual site

There are several ways in which the position of the virtual site can be defined. In the current 2BPA model, the side-chain virtual bead positions are defined by the `3fd` scheme, while the hydrogen bonding bead positions are defined by the `3out` scheme, which has more parameters. Therefore, there is more freedom in the placement of the beads than within the `3fd` scheme. In the `3fd` scheme, the position of the virtual bead ($\mathbf{r}_{\mathrm{vs}}$) is defined by

$$\mathbf{r}_{\mathrm{vs}} = \mathbf{r}_i + b\frac{\mathbf{r}_{i,i-1,i+1}}{|\mathbf{r}_{i,i-1,i+1}|}, \qquad \mathbf{r}_{i,i-1,i+1} = \mathbf{r}_{i,i-1} + a\mathbf{r}_{i-1,i+1}, \tag{2}$$

where $\mathbf{r}_i$ is the position of the connected massive backbone bead, $\mathbf{r}_{i,i-1}$ is the vector from the current massive bead ($\mathbf{r}_i$) to the previous massive bead ($\mathbf{r}_{i-1}$), $\mathbf{r}_{i-1,i+1}$ is the vector from the previous massive bead to the next massive bead ($\mathbf{r}_{i+1}$), $a$ determines the in-plane angle of the virtual bead, and $|b|$ is absolute distance between the current massive bead and the virtual site (see Supplementary Fig.2) and has units of length. As can be seen from Equation 2, the virtual site is always placed in the plane formed by the three massive beads that define its position. The geometry of the `3fd` scheme is further clarified in Supplementary Fig.2.

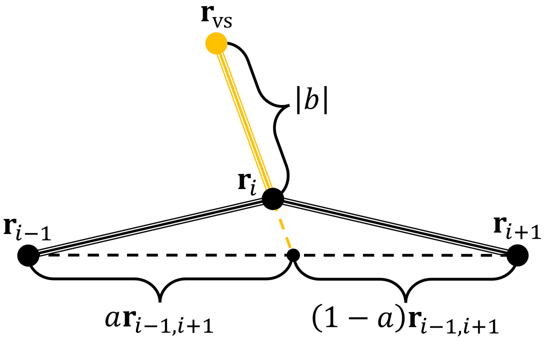


**Supplementary Figure 2: `3fd` virtual site scheme.** The `3fd` scheme for determining the position of the virtual site ($\mathbf{r}_{\mathrm{vs}}$). Here, $a > 0.5$ and $b < 0$.

### Hydrogen bonding locations: `3out` virtual sites

To model the two hydrogen bonding bead (HBO and HBH from Fig. 1) position, `3out` virtual site scheme was chosen. In the `3out` scheme, the position of the virtual site is defined by

$$\mathbf{r}_{\mathrm{vs}} = \mathbf{r}_i + \bar{a}\mathbf{r}_{i,i-1} + \bar{b}\mathbf{r}_{i,i+1} + c\frac{\mathbf{r}_{i,i-1} \times \mathbf{r}_{i,i+1}}{|\mathbf{r}_{i,i-1} \times \mathbf{r}_{i,i+1}|}, \tag{3}$$

where $\mathbf{r}_{i,i-1}$ is the vector between the current massive bead and the previous massive bead, $\mathbf{r}_{i,i+1}$ is the vector between the current massive bead and the next massive bead, and $a$, $b$ and $c$ are parameters that define the position of the virtual site position. Here parameter $c$ has units of length. The geometry of this virtual site type, along with the definitions of $\bar{a}$, $\bar{b}$ and $c$, are clarified in Supplementary Fig. 3. Using this scheme, the virtual bead can also be placed out of the plane of the three massive beads. In the 2BPA model, hydrogen beads use $\bar{a} = 0.333$, $\bar{b} = 0$, and $c = 0.238$, while oxygen beads use $\bar{a} = 0.667$, $\bar{b} = 0$, and $c = -0.238$ For the terminal residues of a protein, there do not exist two neighbouring massive backbone beads. Therefore, the virtual sites are constructed from the second-to-last beads with altered parameters.

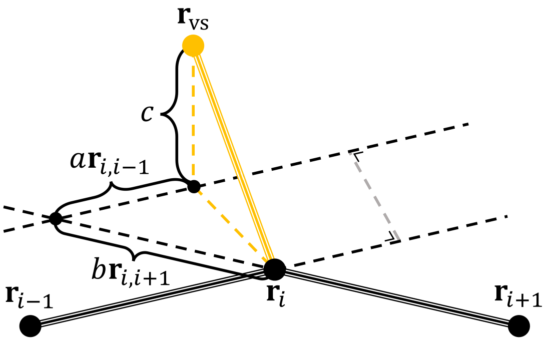


**Supplementary Figure 3:** `3out` **virtual site scheme.** The `3out` scheme for determining the position of the virtual site ($\mathbf{r}_{\text{vs}}$).

**Supplementary Table 1: Secondary structure residues of DNAJB6b.** Ranges of residues containing secondary structure in the DNAJB6b. For each region, the name of the secondary structure, helix (H) or β-sheet (E) is given, along with the domain it resides in.

| Name | Domain | Structure | Residues |
|---|---|---|---|
| H1 | J | α-Helix | [Y4, L8] |
| H2 | J | α-Helix | [P16, W30] |
| H2c | J | 3-10 Helix | [P32, K34] |
| H3 | J | α-Helix | [K39, S57] |
| H4 | J | α-Helix | [A59, N74] |
| H5 | $G/F_1$ | α-Helix | [P96, F104] |
| H6 | $G/F_2$ | α-Helix | [F112, E120] |
| H7 | S/T | α-Helix | [T137, F144] |
| H8 | S/T | α-Helix | [S156, F164] |
| E1 | CTD | β-Strand | [S190, V198] |
| E2 | CTD | β-Strand | [R201, E210] |
| E3 | CTD | β-Strand | [E214, E220] |
| E4 | CTD | β-Strand | [Q223, I229] |
| E5 | CTD | β-Strand | [K232, Q234] |

**Supplementary Table 2: `3fd` virtual bead parameters for DNAJB6b.** Side-chain virtual bead parameters of DNAJB6b, split up into the different secondary structure types, obtained from all-atom MD simulations. The modes obtained from the combined distributions of the four secondary structures is given under the `total` columns.

| | `3fd` $a$ | | | | | `3fd` $\|b\|$ (nm) | | | | |
|---|---|---|---|---|---|---|---|---|---|---|
| Residue | Total | Helix | Sheet | Turn | Coil | Total | Helix | Sheet | Turn | Coil |
| ALA (A) | 0.363 | 0.363 | -† | 0.449 | 0.484 | 0.162 | 0.162 | -† | 0.161 | 0.162 |
| ARG (R) | 0.514 | 0.647 | 0.480 | 0.549 | 0.497 | 0.422 | 0.422 | 0.441 | 0.422 | 0.426 |
| ASN (N) | 0.596 | 0.783 | 0.572 | 0.888 | 0.586 | 0.256 | 0.256 | 0.257 | 0.256 | 0.255 |
| ASP (D) | 0.580 | 0.682 | 0.316 | 0.692 | 0.566 | 0.255 | 0.255 | 0.254 | 0.255 | 0.255 |
| GLN (Q) | 0.565 | 0.599 | 0.587 | 0.547 | 0.565 | 0.267 | 0.345 | 0.267 | 0.266 | 0.267 |
| GLU (E) | 0.544 | 0.582 | 0.546 | 0.534 | 0.523 | 0.343 | 0.342 | 0.343 | 0.343 | 0.342 |
| HIS (H) | 0.205 | 0.866 | 0.579 | 0.205 | 0.192 | 0.320 | 0.319 | 0.321 | 0.320 | 0.321 |
| ILE (I) | 0.534 | 0.514 | 0.534 | 0.517 | 0.517 | 0.245 | 0.246 | 0.245 | 0.210 | 0.245 |
| LEU (L) | 0.546 | 0.644 | 0.361 | 0.747 | 0.546 | 0.271 | 0.270 | 0.272 | 0.271 | 0.270 |
| LYS (K) | 0.613 | 0.634 | 0.560 | 0.817 | 0.562 | 0.377 | 0.375 | 0.377 | 0.332 | 0.377 |
| MET (M) | 0.544 | 0.691 | 0.538 | 0.656 | 0.556 | 0.301 | 0.300 | 0.305 | 0.304 | 0.301 |
| PHE (F) | 0.603 | 0.136 | 0.598 | 0.609 | 0.606 | 0.344 | 0.345 | 0.344 | 0.344 | 0.343 |
| PRO (P) | 0.769 | 1.250 | 0.741 | 1.266 | 0.778 | 0.192 | 0.192 | 0.191 | 0.193 | 0.193 |
| SER (S) | 0.432 | 0.255 | 0.432 | 0.355 | 0.364 | 0.201 | 0.200 | 0.201 | 0.200 | 0.201 |
| THR (T) | 0.641 | 0.449 | 0.469 | 0.451 | 0.439 | 0.199 | 0.199 | 0.199 | 0.200 | 0.200 |
| TRP (W) | 0.812 | 0.812 | -† | 0.879 | 0.821 | 0.370 | 0.370 | -† | 0.366 | 0.368 |
| TYR (Y) | 0.779 | 0.779 | -† | 0.823 | 0.918 | 0.388 | 0.388 | -† | 0.389 | 0.388 |
| VAL (V) | 0.491 | 0.407 | 0.491 | 0.512 | 0.514 | 0.201 | 0.200 | 0.199 | 0.200 | 0.200 |
| CYS (C) | - | - | - | - | - | - | - | - | - | - |

† These residues were never in a sheet conformation.

#### 3fd scheme parameter histograms

The following figures (4–21) show the obtained histograms of the `3fd` virtual bead parameters for all the different amino acid types. The modes given in all of these plots are summarized in Supplementary Table 2. Each figure shows the obtained `3fd` $a$ and $|b|$ parameters, split up into different secondary structures. Furthermore, Supplementary Fig.22 shows the `3fd` $a$ and $|b|$ parameters obtained for polyQ from a CHARMM36m simulation of 20 Q48 molecules. The modes for this plot are significantly different from those obtained from Fig. 3, which were obtained from an AAMD simulation of a Q48 amyloid fibril using CHARMM36m forcefield.

In the following supplementary figures (4–21), each row shows the probability density (PD) distribution for a different secondary structure type. The most probable values are given in each subplot as the mode, highlighted with the dashed lines with `3fd` parameter $|b|$ in angstroms (Å).

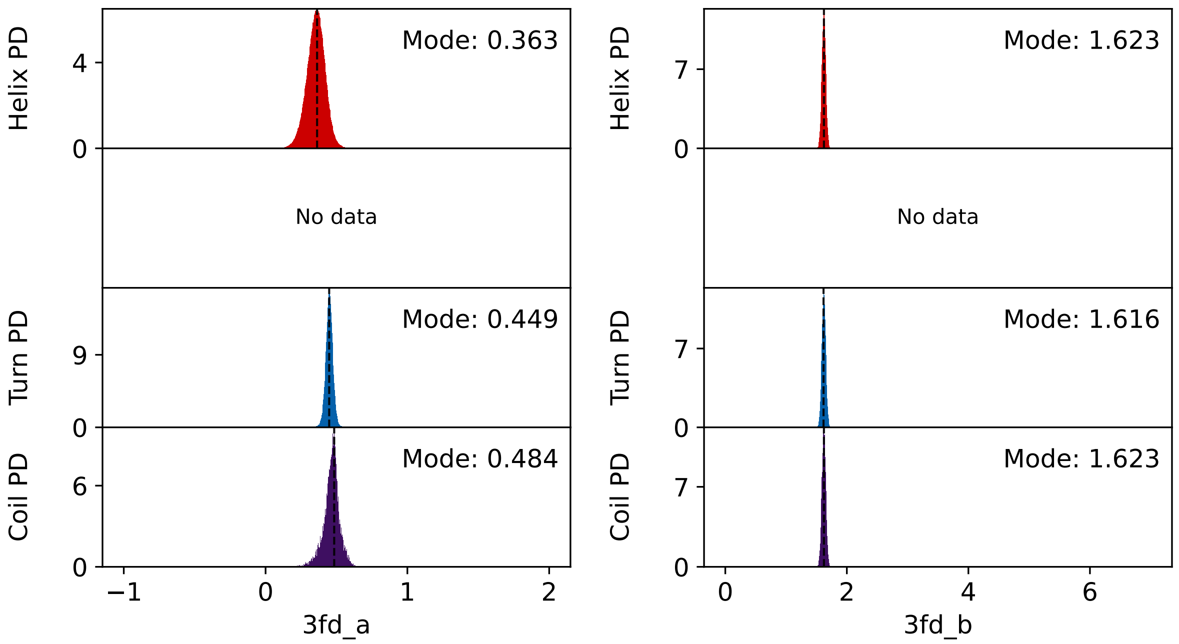


**Supplementary Figure 4: `3fd` virtual bead parameters for the amino acid: alanine.** Probability density (PD) distributions of the `3fd` scheme parameters $a$ and $|b|$ for alanine (Ala, A).

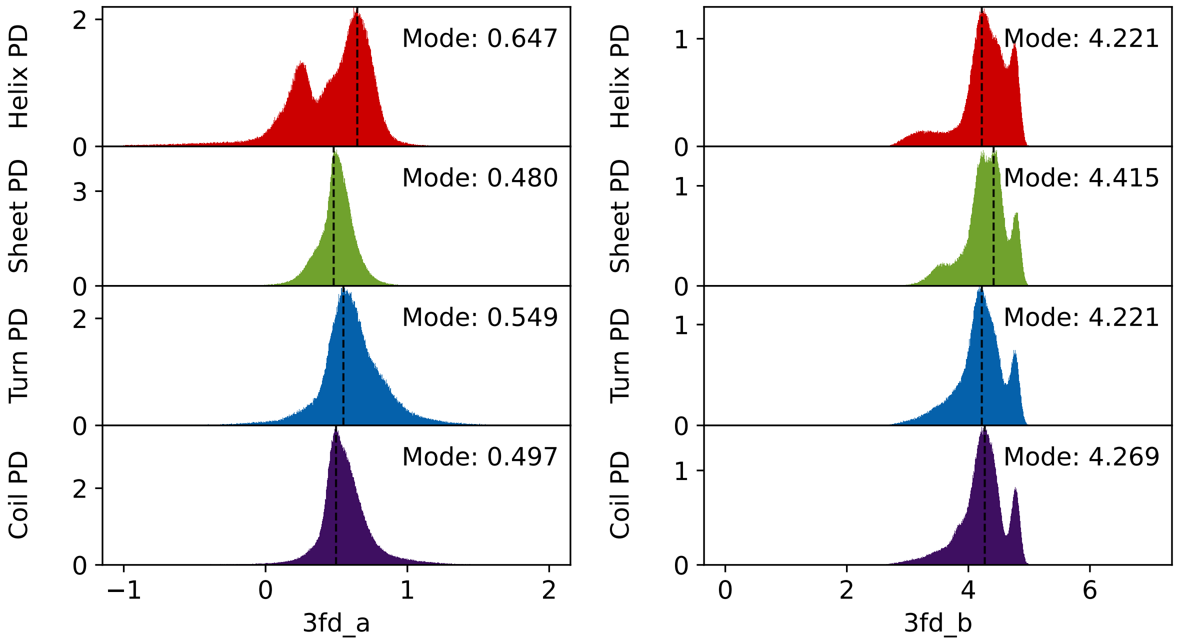


**Supplementary Figure 5: `3fd` virtual bead parameters for the amino acid: arginine.** Probability density (PD) distributions of the `3fd` scheme parameters $a$ and $|b|$ for arginine (Arg, R).

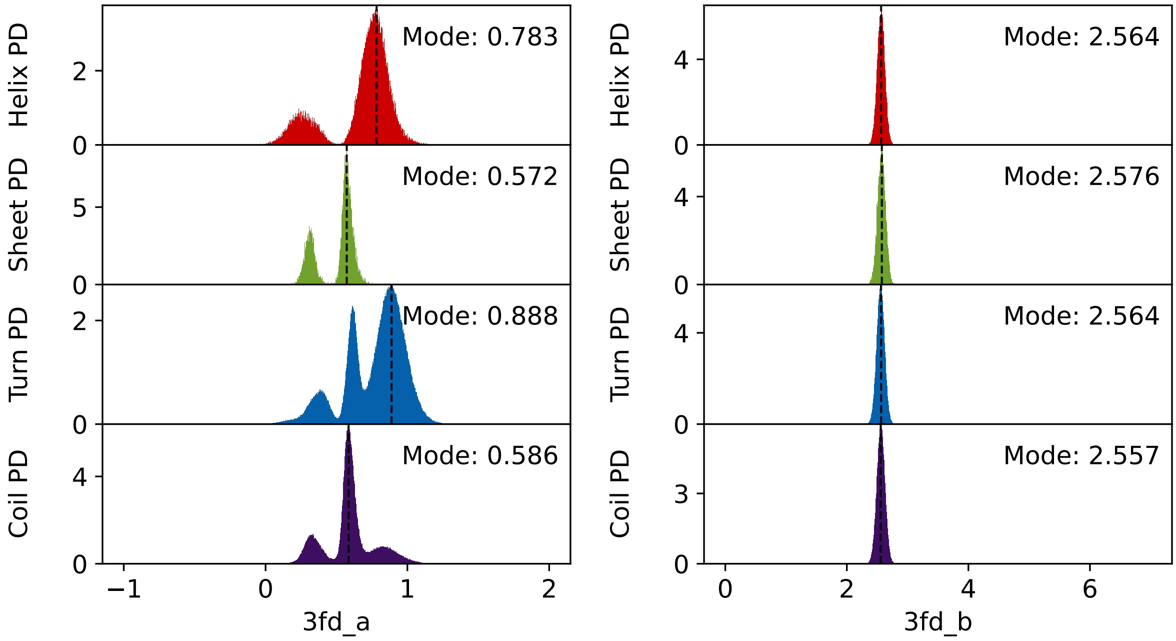


**Supplementary Figure 6: `3fd` virtual bead parameters for the amino acid: aspargine.** Probability density (PD) distributions of the `3fd` scheme parameters $a$ and $|b|$ for asparagine (Asn, N).

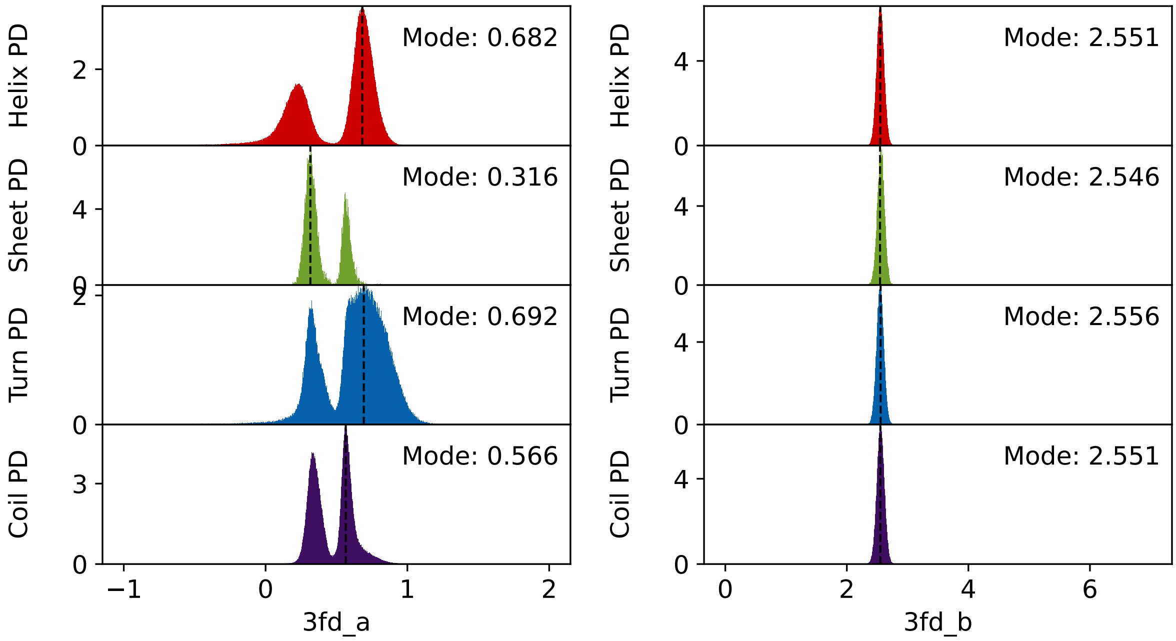


**Supplementary Figure 7: `3fd` virtual bead parameters for the amino acid: aspartic acid.** Probability density (PD) distributions of the `3fd` scheme parameters $a$ and $|b|$ for aspartic acid (Asp, D).

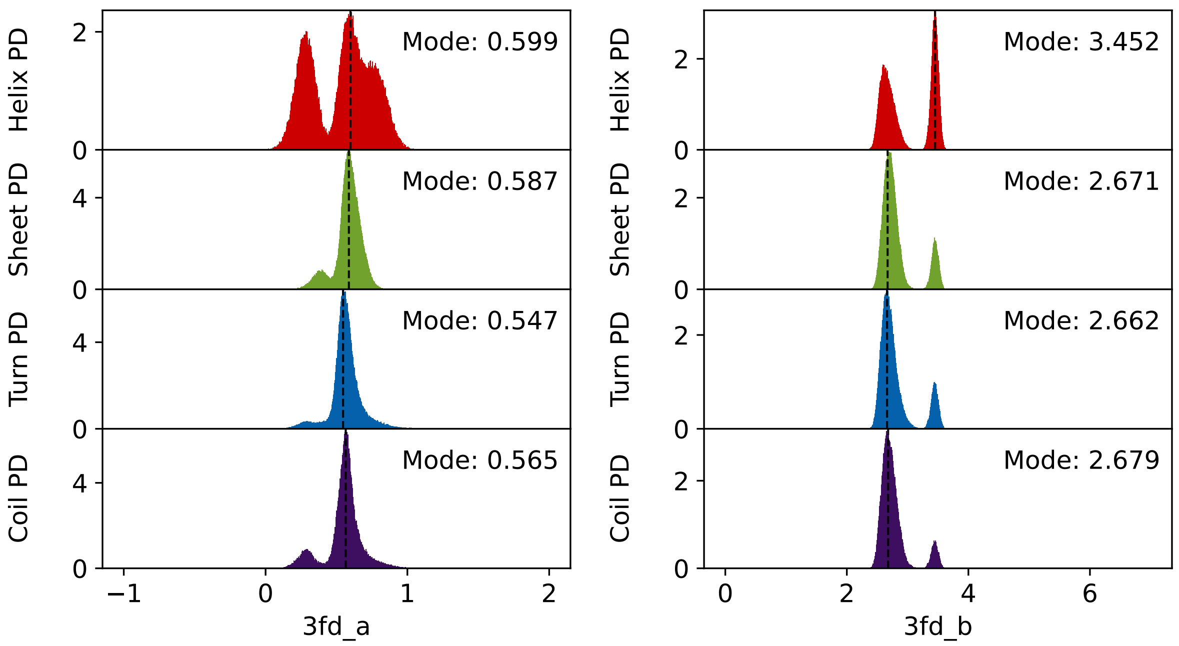


**Supplementary Figure 8: `3fd` virtual bead parameters for the amino acid: glutamine.** Probability density (PD) distributions of the `3fd` scheme parameters $a$ and $|b|$ for glutamine (Gln, Q).

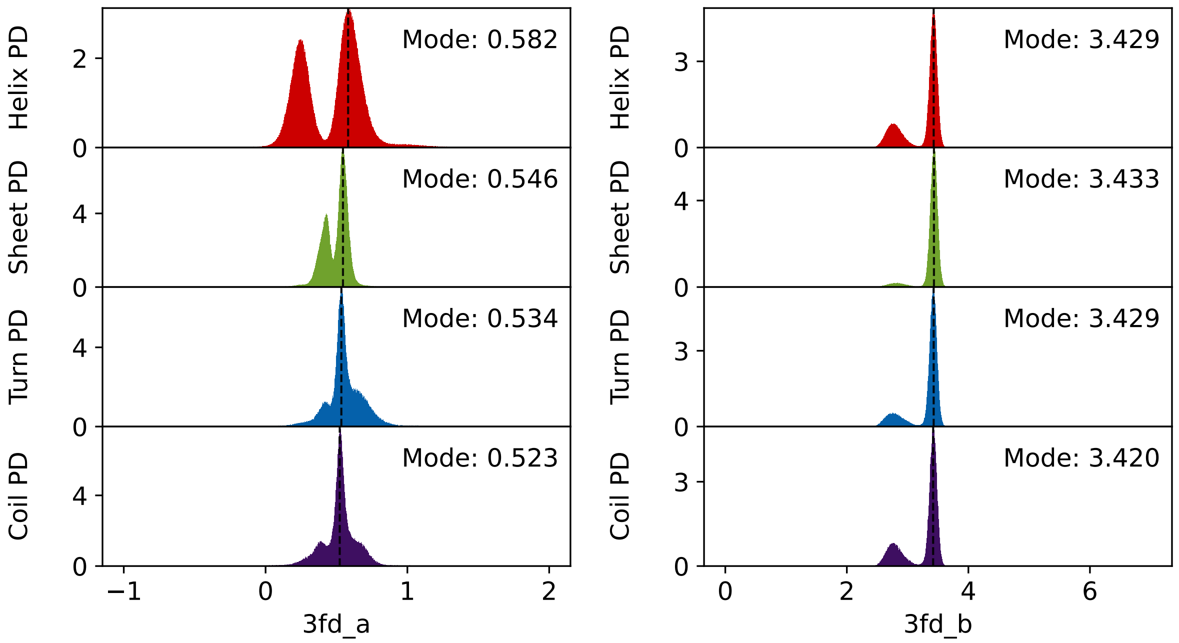


**Supplementary Figure 9: `3fd` virtual bead parameters for the amino acid: glutamic acid.** Probability density (PD) distributions of the `3fd` scheme parameters $a$ and $|b|$ for glutamic acid (Glu, E).

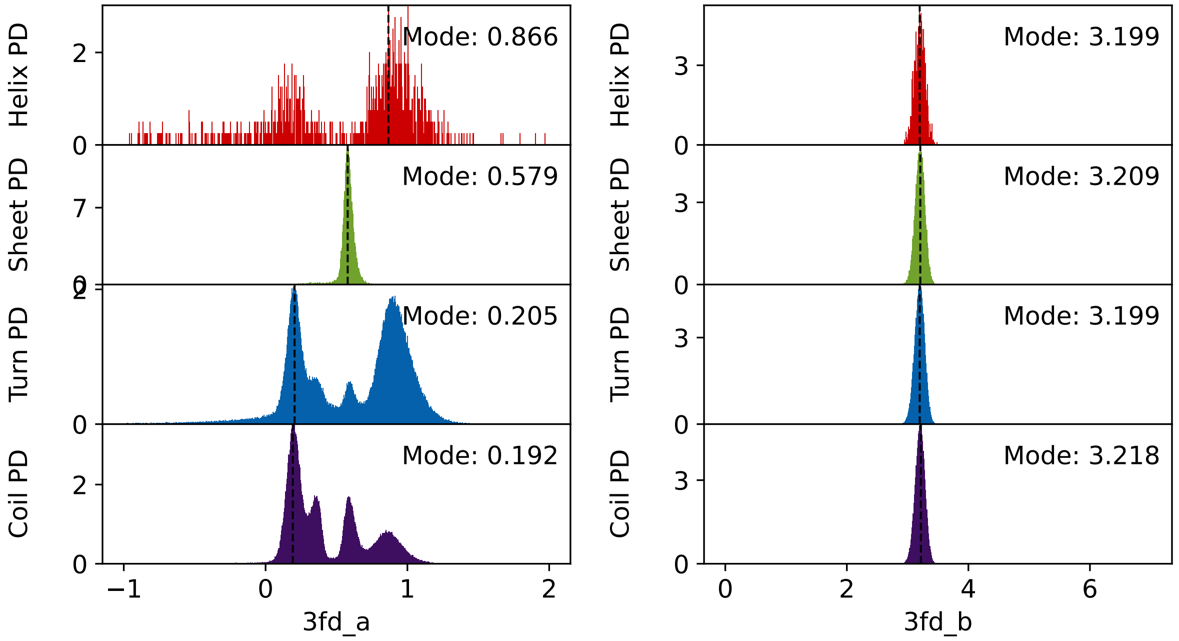


**Supplementary Figure 10: `3fd` virtual bead parameters for the amino acid: histidine.** Probability density (PD) distributions of the `3fd` scheme parameters $a$ and $|b|$ for histidine (His, H).

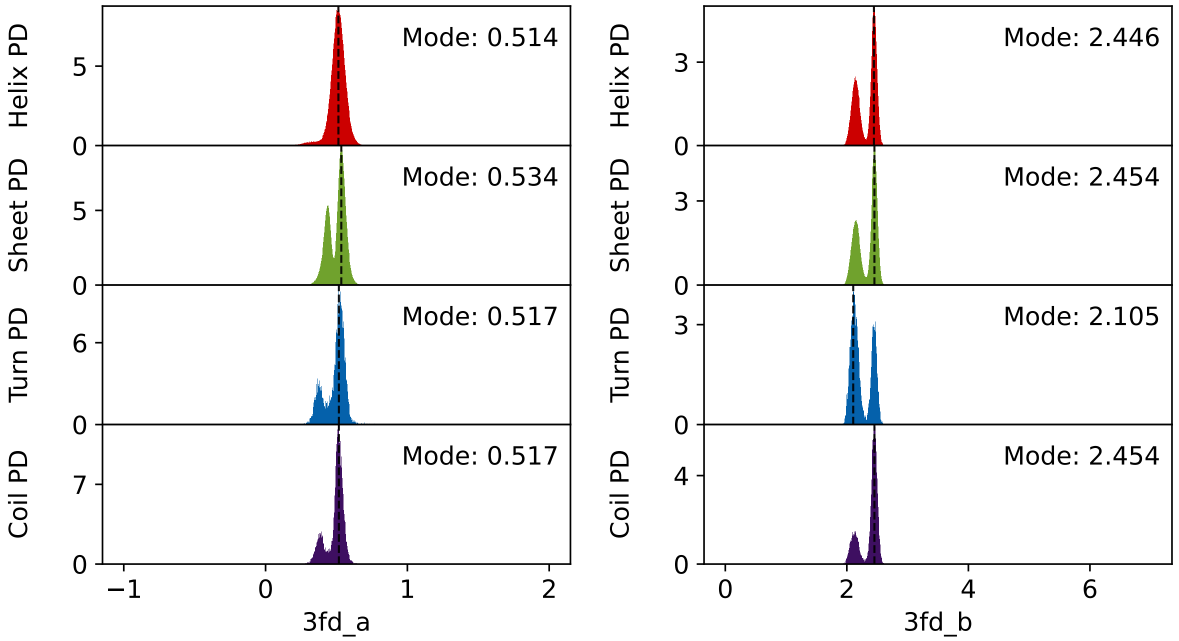


**Supplementary Figure 11: `3fd` virtual bead parameters for the amino acid: isoleucine.** Probability density (PD) distributions of the `3fd` scheme parameters $a$ and $|b|$ for isoleucine (Ile, I).

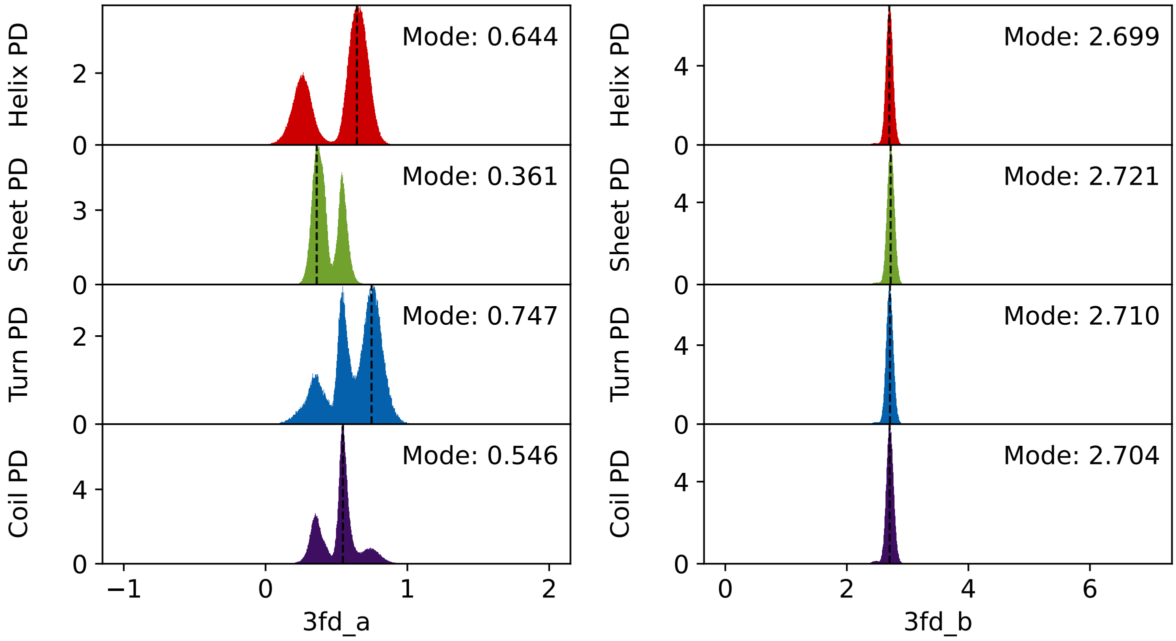


**Supplementary Figure 12: `3fd` virtual bead parameters for the amino acid: leucine.** Probability density (PD) distributions of the `3fd` scheme parameters $a$ and $|b|$ for leucine (Leu, L).

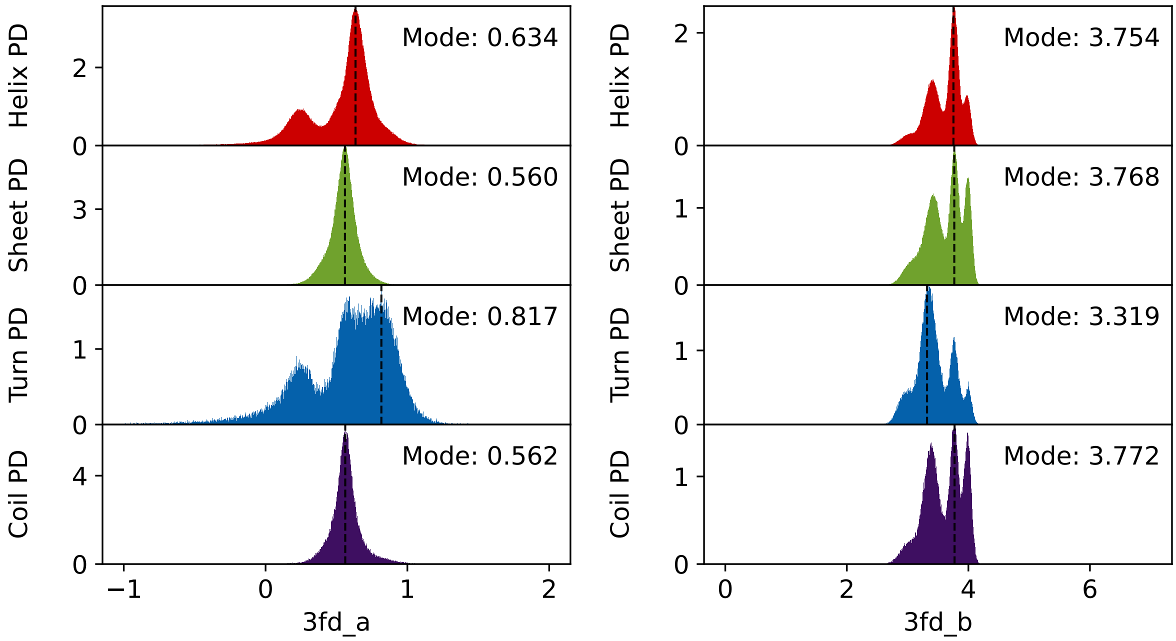


**Supplementary Figure 13: `3fd` virtual bead parameters for the amino acid: lysine.** Probability density (PD) distributions of the `3fd` scheme parameters $a$ and $|b|$ for lysine (Lys, K).

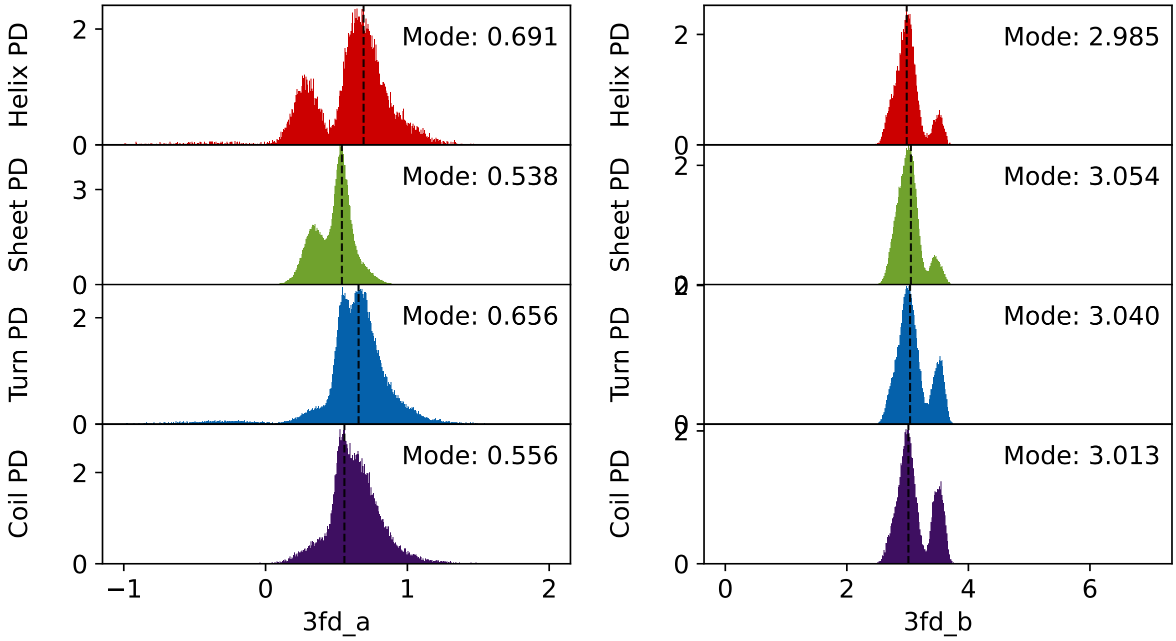


**Supplementary Figure 14: `3fd` virtual bead parameters for the amino acid: methionine.** Probability density (PD) distributions of the `3fd` scheme parameters $a$ and $|b|$ for methionine (Met, M).

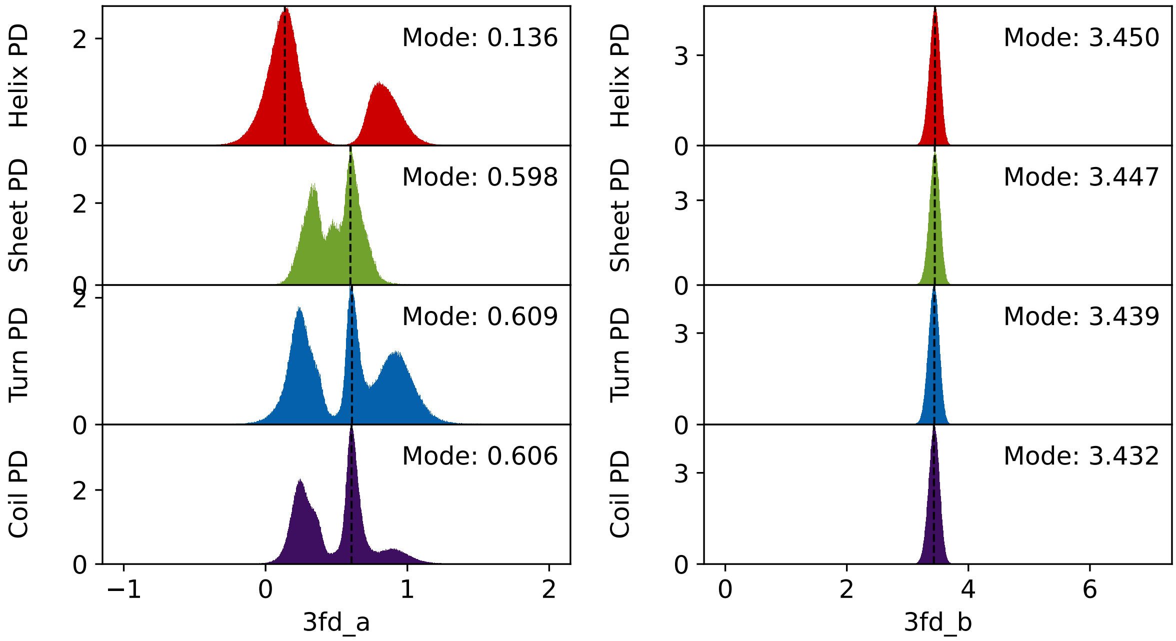


**Supplementary Figure 15: `3fd` virtual bead parameters for the amino acid: phenylalanine.** Probability density (PD) distributions of the `3fd` scheme parameters $a$ and $|b|$ for phenylalanine (Phe, F).

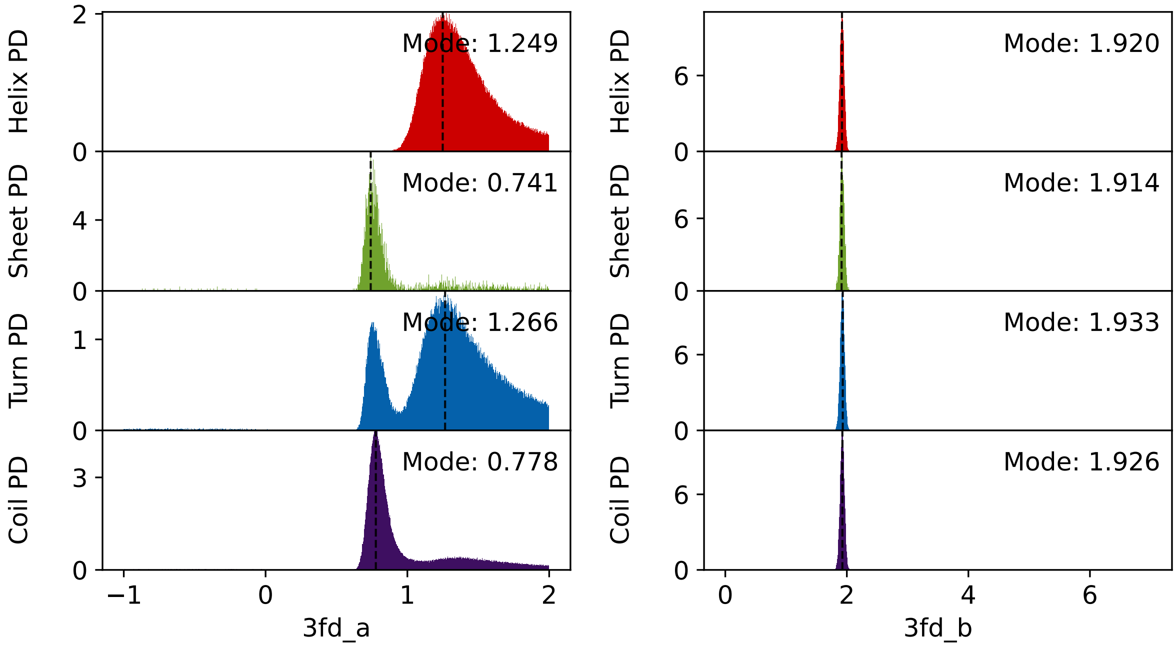


**Supplementary Figure 16: `3fd` virtual bead parameters for the amino acid: proline.** Probability density (PD) distributions of the `3fd` scheme parameters $a$ and $|b|$ for proline (Pro, P).

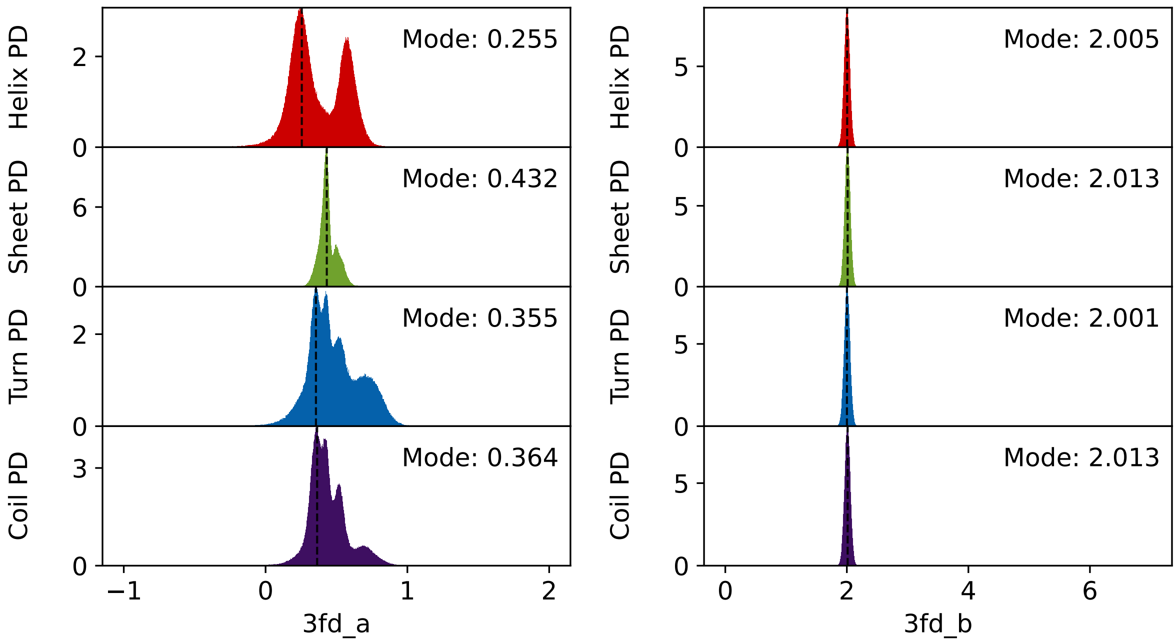


**Supplementary Figure 17: `3fd` virtual bead parameters for the amino acid: serine.** Probability density (PD) distributions of the `3fd` scheme parameters $a$ and $|b|$ for serine (Ser, S).

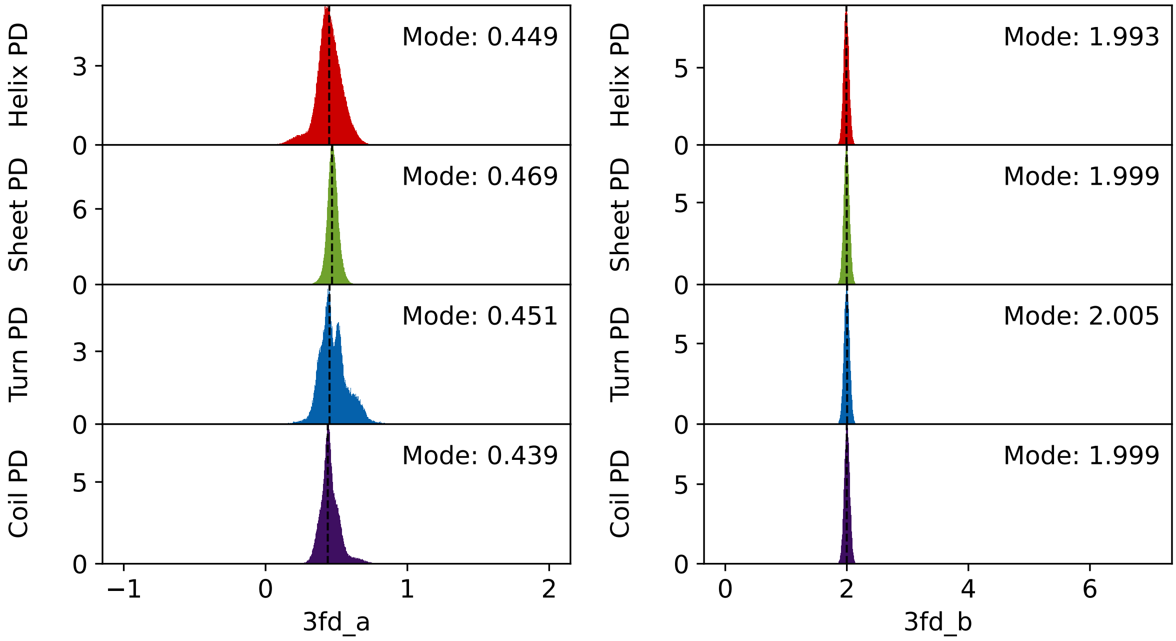


**Supplementary Figure 18: `3fd` virtual bead parameters for the amino acid: threonine.** Probability density (PD) distributions of the `3fd` scheme parameters $a$ and $|b|$ for threonine (Thr, T).

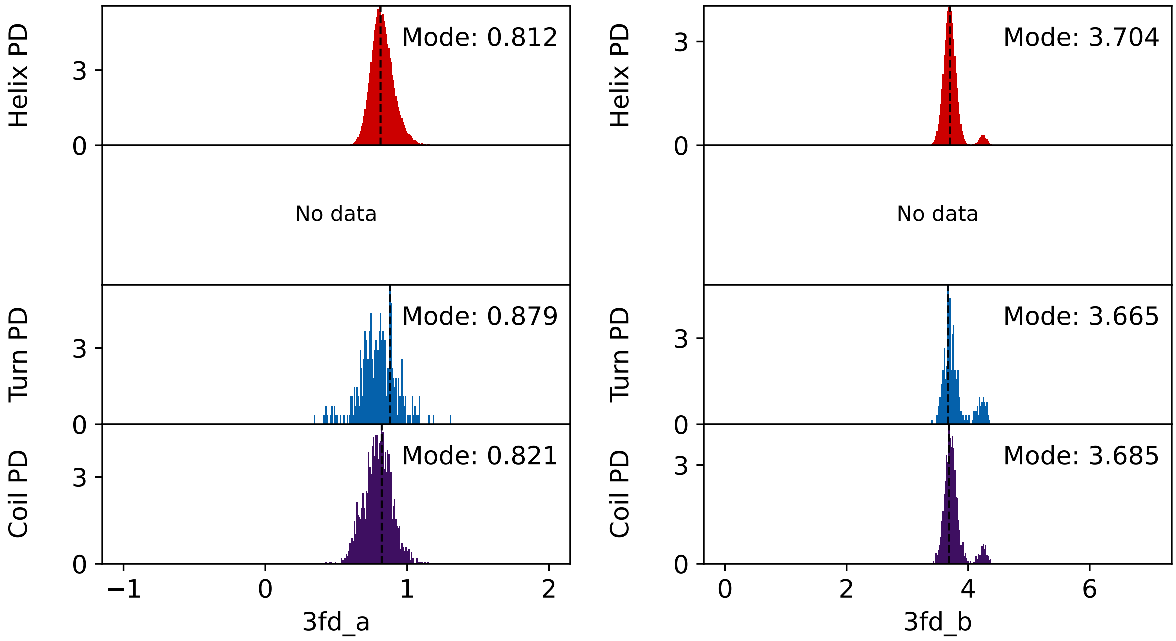


**Supplementary Figure 19: `3fd` virtual bead parameters for the amino acid: tryptophan.** Probability density (PD) distributions of the `3fd` scheme parameters $a$ and $|b|$ for tryptophan (Trp, W).

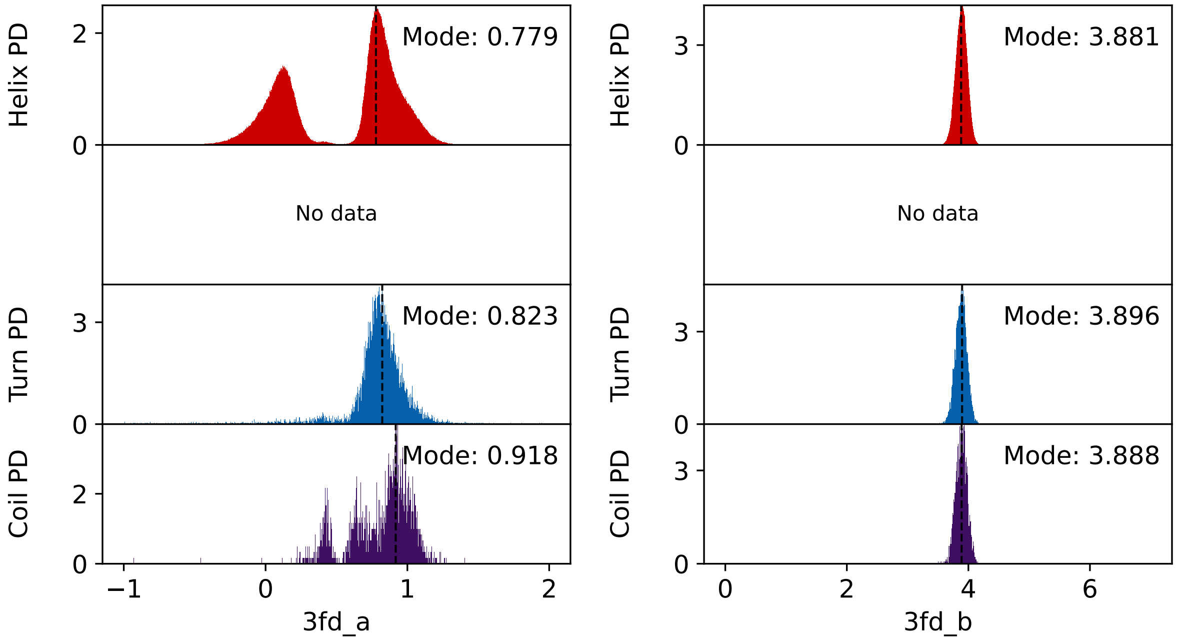


**Supplementary Figure 20: `3fd` virtual bead parameters for the amino acid: tyrosine.** Probability density (PD) distributions of the `3fd` scheme parameters $a$ and $|b|$ for tyrosine (Tyr, Y).

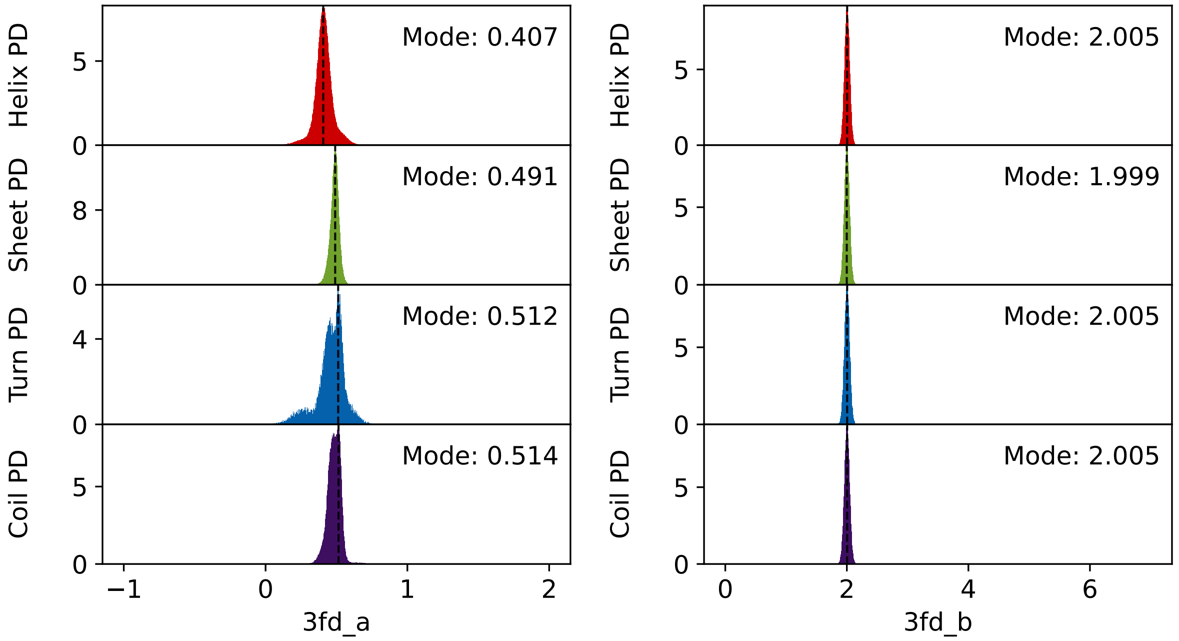


**Supplementary Figure 21: `3fd` virtual bead parameters for the amino acid: valine.** Probability density (PD) distributions of the `3fd` scheme parameters $a$ and $|b|$ for valine (Val, V).

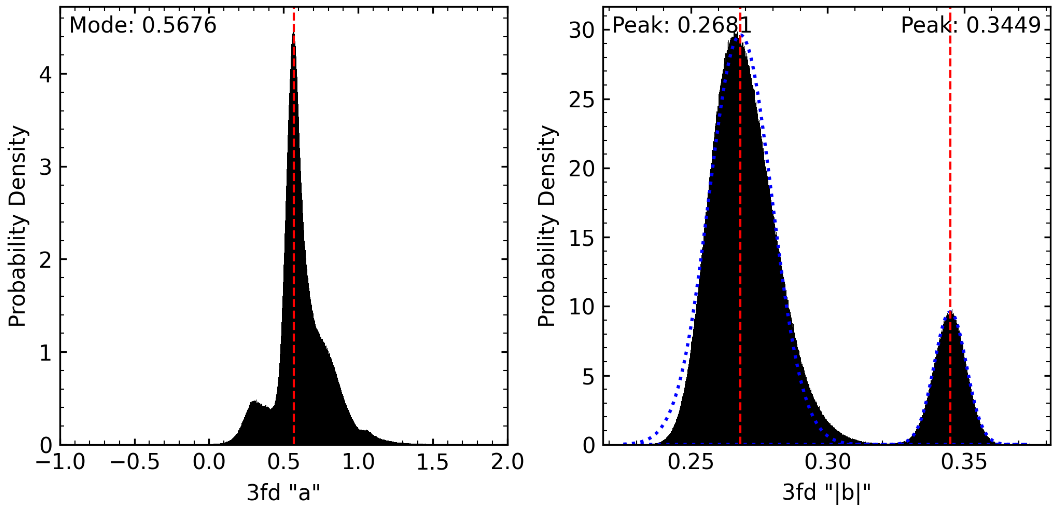


**Supplementary Figure 22: `3fd` virtual bead parameters for the amino acid: glutamine of polyglutamine peptide.** Histograms of 3fd $a$ and 3fd $|b|$. Data was obtained from an all-atom simulations of 20 Q48 molecules. The `3fd` parameter $b$ has units of length.

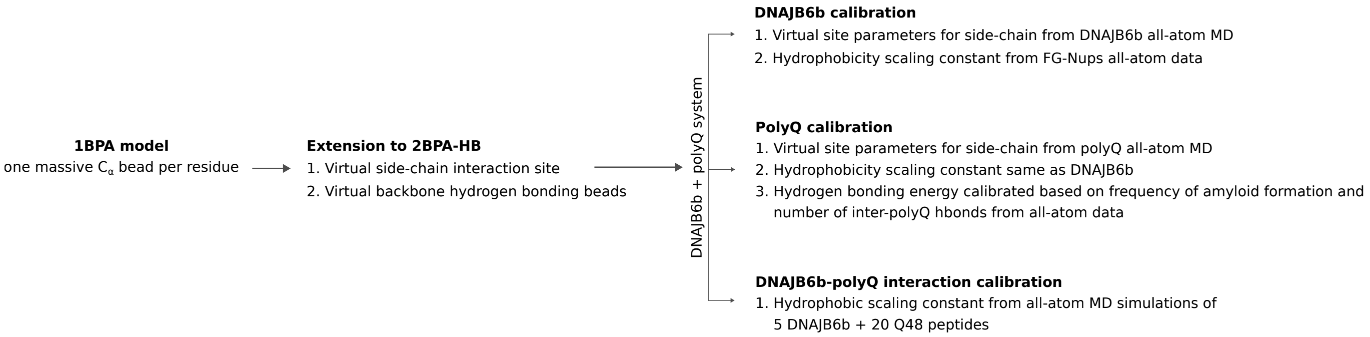


**Supplementary Figure 23: Workflow for the development and calibration of the 2BPA-HB model.** The 1BPA framework, containing one massive $C_\alpha$ bead per residue, was extended by introducing virtual side-chain interaction sites and virtual backbone hydrogen-bonding sites. The model was subsequently calibrated in three stages. DNAJB6b parameters were obtained from all-atom MD data for side-chain virtual-site geometry and from FG-Nup simulations for the hydrophobic scaling constant. PolyQ parameters were obtained from all-atom polyQ simulations, with the hydrogen-bonding energy calibrated using the number of inter-polyQ hydrogen bonds and the frequency of amyloid-like structure formation. Finally, the DNAJB6b–polyQ interaction strength was calibrated using all-atom MD simulations containing five DNAJB6b molecules and 20 Q48 peptides.

**Supplementary Table 3: Comparision of $R_g$ of DNAJB6b.** Radius of gyration of DNAJB6b for all three states, compared to AAMD simulations.

| State | CG $R_g$ [nm] | AA $R_g$ [nm] |
|---|---|---|
| Closed | $2.31 \pm 0.10$ | $2.10 \pm 0.10$ |
| Open | $2.71 \pm 0.27$ | $2.56 \pm 0.25$ |
| Extended | $3.17 \pm 0.39$ | $3.06 \pm 0.53$ |

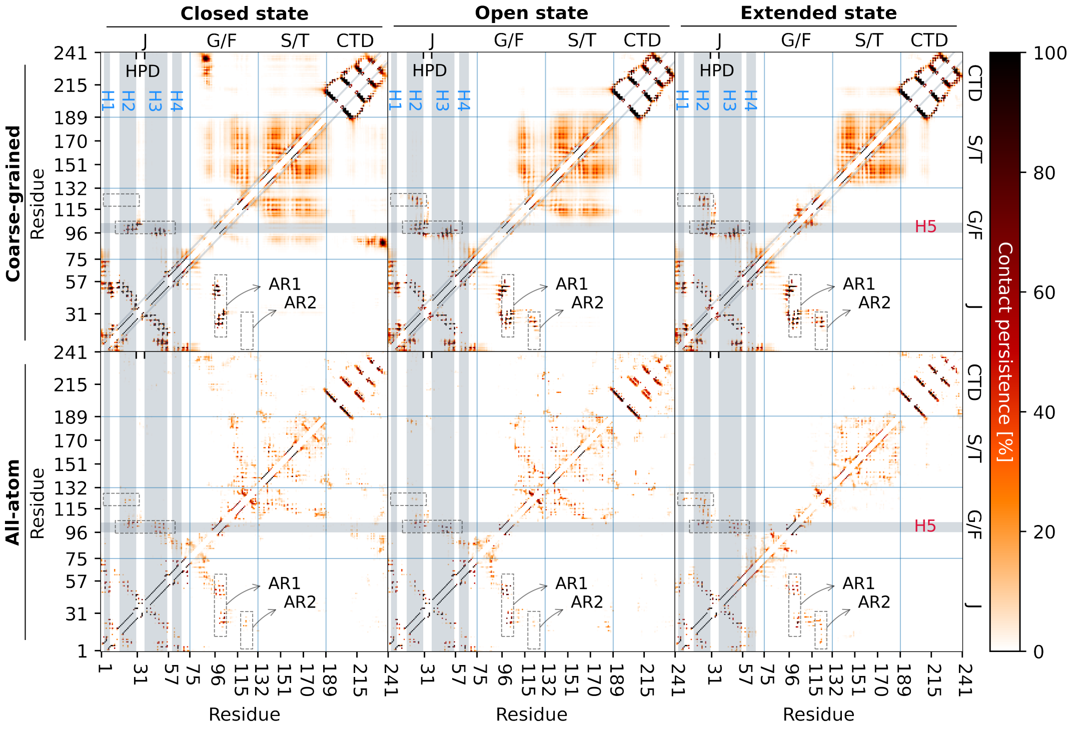


**Supplementary Figure 24: Intramolecular contact maps of DNAJB6b.** Intra-molecular DNAJB6b contact maps, split up into the closed, open, and extended state of DNAJB6b. The top charts show the results from the best CG model, and the bottom charts show the contact maps from the AAMD simulations.

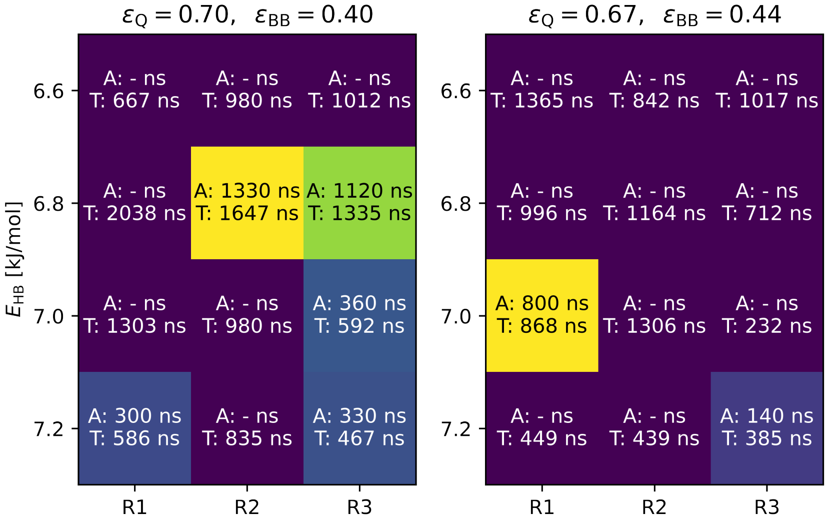


**Supplementary Figure 25: Parametrization of hydrophobic scaling constant and hydrogen-bonding energy of polyglutamine peptide.** The total simulation time (T) and time at which amyloid formation first occurred (A), for different values of side-chain hydrophobicity $\varepsilon_{\mathrm{Q}}$, backbone hydrophobicity $\varepsilon_{\mathrm{BB}}$ and hydrogen bonding energy $E_{\mathrm{HB}}$. The colors show the amyloid formation time, where purple means no amyloid formation occurred at all, blue indicates early amyloid formation, and yellow indicates late amyloid formation. Three replicates were used (R1, R2 and R3).

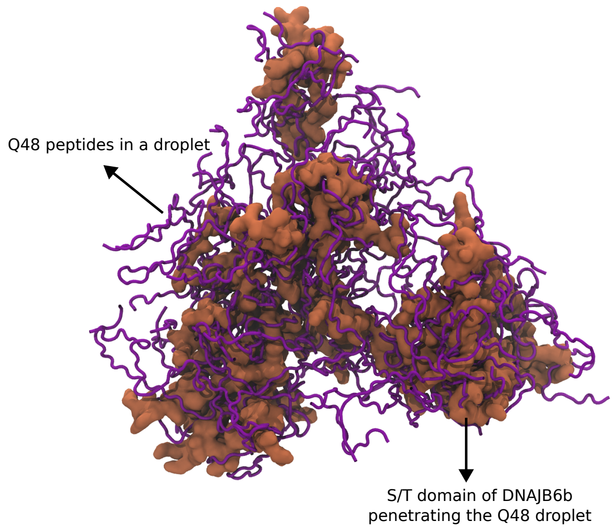


**Supplementary Figure 26: Penetration of the S/T domain of DNAJB6b into a polyQ droplet.** Final snapshot of a CGMD simulation with 52 Q48 peptides and 13 DNAJB6b molecules, initialized from a droplet configuration. For clarity, only the S/T domain of DNAJB6b is shown, illustrating its penetration into the Q48 peptide droplet.

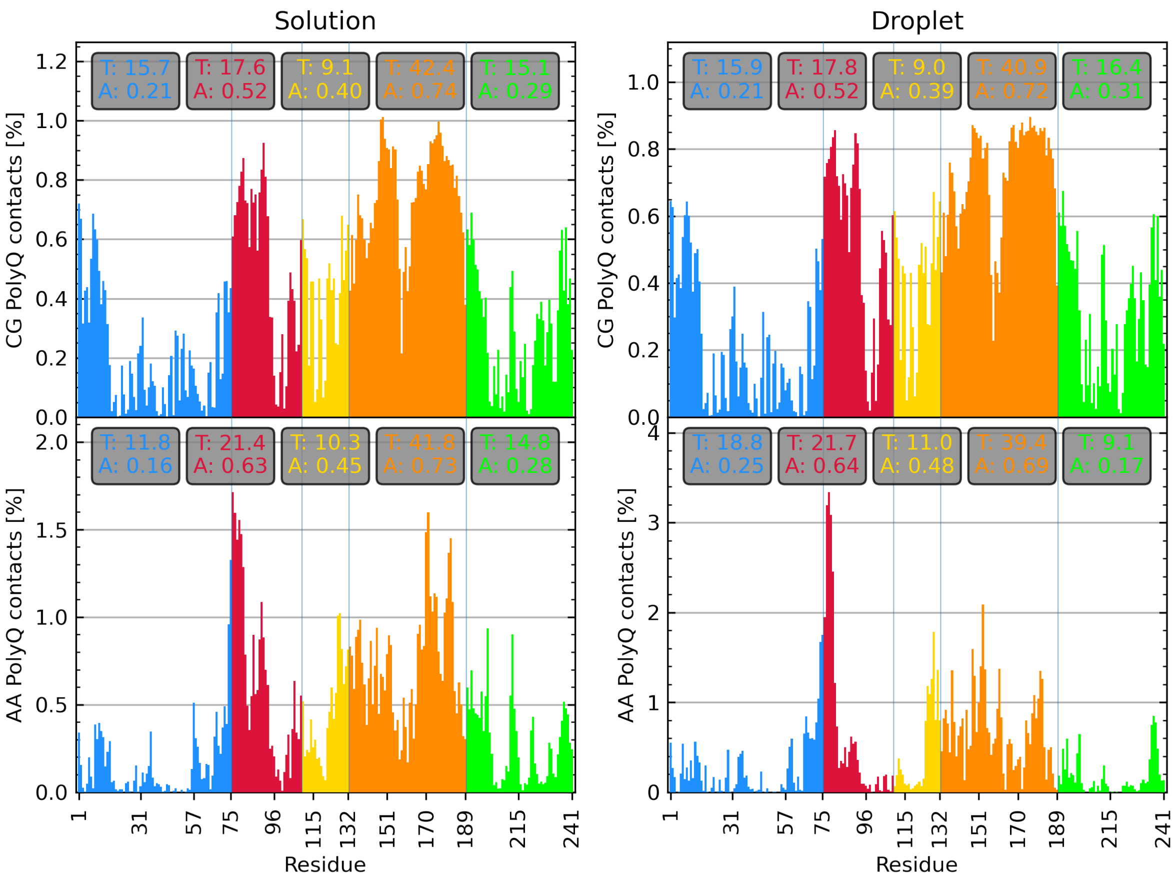


**Supplementary Figure 27: Domain level contacts between the chaperone DNAJB6b and polyglutamine.** Contact bar charts showing which residues of DNAJB6b interact with polyQ. The top graphs show the CG distributions with $\alpha_{\text{JB6-Q}} = 0.13$, while the bottom graphs show the AA results. The left graphs show contacts when starting with a polyQ solution, while for the right graphs the initial configuration was a polyQ droplet. The total percentage per domain (T), as well as the average percentage per domain (A) are also given.

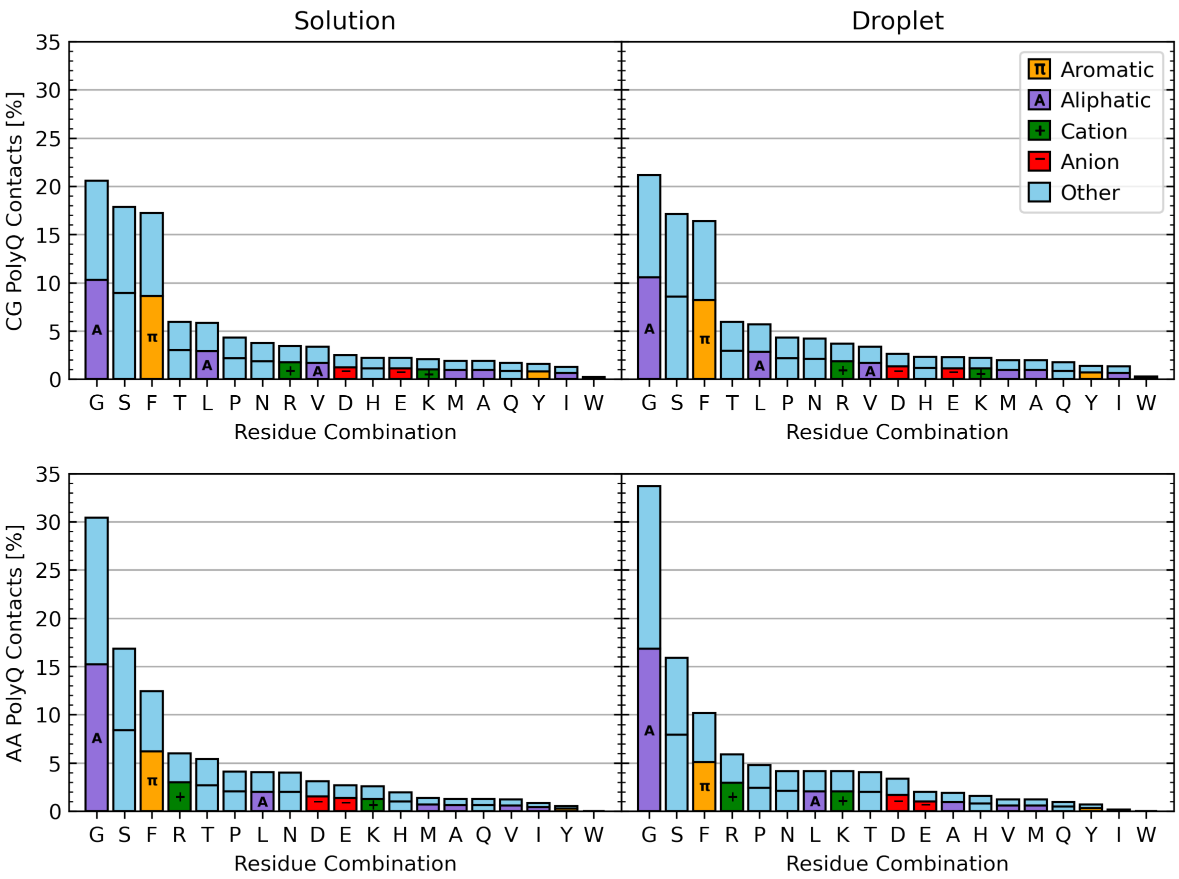


**Supplementary Figure 28: Amino-acid level contacts between the chaperone DNAJB6b and polyglutamine.** Residue pair bar charts showing which amino acid types of DNAJB6b interact with polyQ. The top graphs show the CG distributions with $\alpha_{\text{JB6-Q}} = 0.13$, while the bottom graphs show the AA results. The left graphs show contacts when starting with a polyQ solution, whilst for the right graphs the initial configuration was a polyQ droplet.

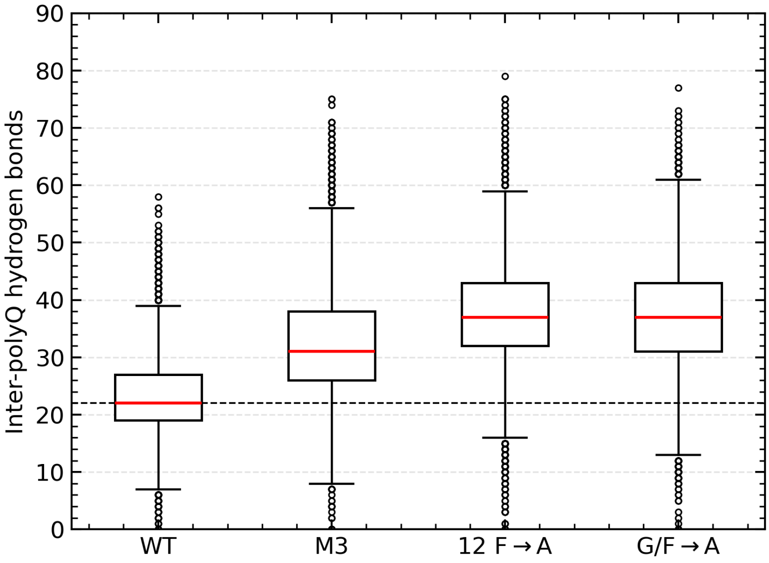


**Supplementary Figure 29: Effect of DNAJB6b mutations on inter-polyQ hydrogen bonding.** Box plots show the number of intermolecular hydrogen bonds between Q48 molecules in simulations containing 52 Q48 and 13 open-state DNAJB6b molecules. Wild-type DNAJB6b is compared with the M3 mutant, in which 12 S and six T residues in the S/T-rich region were substituted by A; the 12 F→A mutant, in which all F residues in the S/T-rich region were substituted by A; and the G/F→A mutant, in which all G and F residues in the G/$F_1$-rich region were substituted by A. Data from the final quarter of three replica simulations of at least 1 µs were combined for each condition. Red lines indicate the median, boxes represent the interquartile range, and the horizontal dashed line indicates the wild-type median.

### RMSF

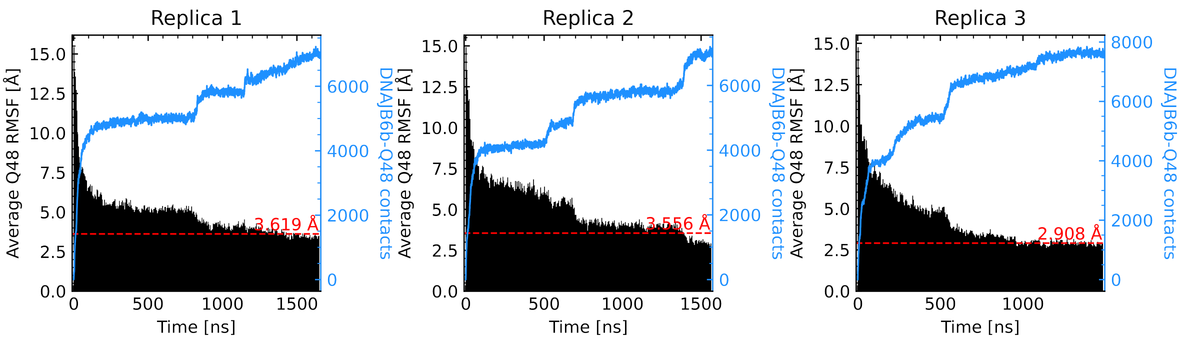


**Supplementary Figure 30: Effect of DNAJB6b interactions on the RMSF of polyglutamine peptides starting from a droplet state.** The average RMSF of 52 Q48 molecules and the number of contacts with the 13 DNAJB6b molecules over time, starting with a polyQ droplet for three replicas. The average RMSF of the last quarter of each simulation is given by the red dashed line.

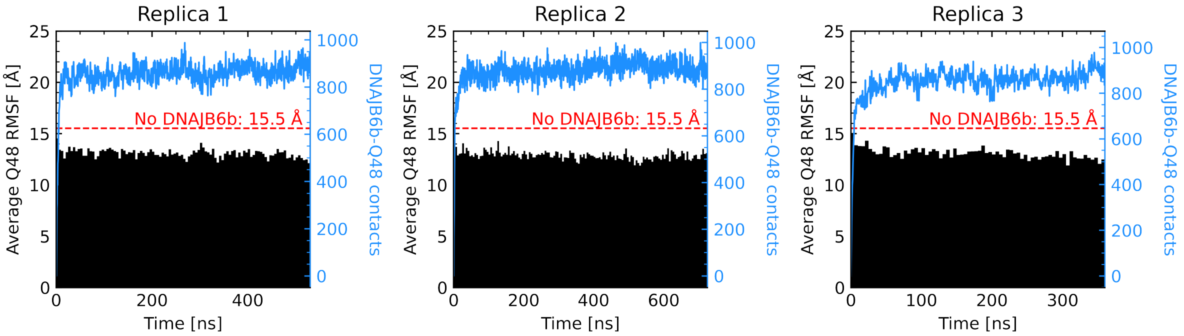


**Supplementary Figure 31: Effect of a single DNAJB6b interactions on the RMSF of polyglutamine peptides starting from a droplet state.** The average RMSF of 52 Q48 molecules and the number of contacts with a single DNAJB6b molecule over time, starting with a polyQ droplet for three replicas. The average RMSF of simulations without DNAJB6b is given by the red dashed line.

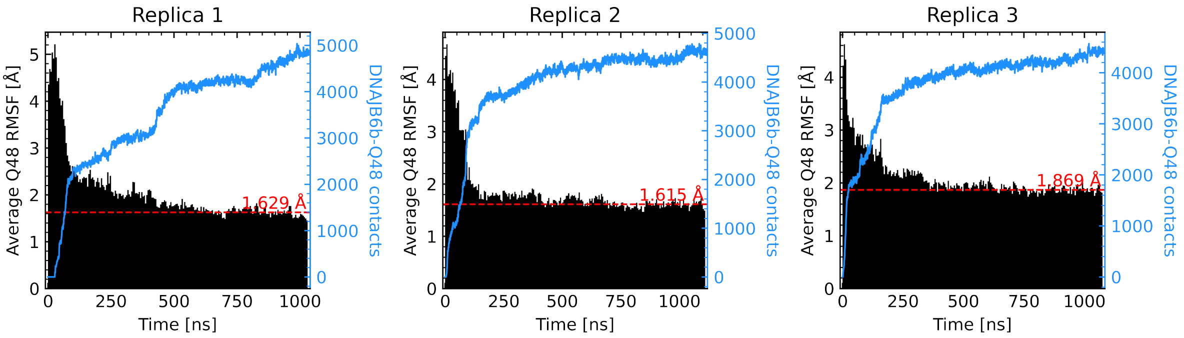


**Supplementary Figure 32: Effect of DNAJB6b interactions on the RMSF of polyglutamine peptides starting from an amyloid state.** The average RMSF of 52 Q48 molecules and the number of contacts with 13 DNAJB6b molecules over time, starting with a polyQ amyloid for three replicas. The average RMSF of the last quarter of each simulation is given by the red dashed line.

**Supplementary Table 4: Conformational sampling of DNAJB6b.** The final composition of states of DNAJB6b in the simulations starting with mixed, all-closed, all-open, and all-extended DNAJB6b. The percentages of states are given for the final 50 ns of the simulations.

| | | Closed state [%] | | | Open state [%] | | | Extended state [%] | | |
|---|---|---|---|---|---|---|---|---|---|---|
| DNAJB6b† | Q48† | R1 | R2 | R3 | R1 | R2 | R3 | R1 | R2 | R3 |
| Mixed | Solution | 45 | 46 | 39 | 15 | 12 | 4 | 39 | 42 | 57 |
| | Droplet | 40 | 39 | 41 | 12 | 11 | 8 | 47 | 50 | 51 |
| | Amyloid | 39 | 39 | 38 | 14 | 8 | 9 | 47 | 54 | 53 |
| All closed | Solution | 99 | 99 | 100 | 1 | 1 | 0 | 0 | 0 | 0 |
| | Droplet | 99 | 99 | 100 | 1 | 1 | 0 | 0 | 0 | 0 |
| | Amyloid | 98 | 100 | 100 | 2 | 0 | 0 | 0 | 0 | 0 |
| All open | Solution | 2 | 8 | 1 | 14 | 16 | 14 | 85 | 76 | 86 |
| | Droplet | 2 | 0 | 2 | 22 | 14 | 19 | 77 | 86 | 79 |
| | Amyloid | 1 | 0 | 2 | 23 | 15 | 30 | 77 | 84 | 68 |
| All extended | Solution | 3 | 3 | 1 | 12 | 14 | 19 | 85 | 82 | 81 |
| | Droplet | 0 | 1 | 0 | 14 | 24 | 10 | 86 | 75 | 90 |
| | Amyloid | 0 | 2 | 0 | 23 | 18 | 6 | 77 | 80 | 93 |

† Initial configuration of the molecules.

#### Hydrophobicities of amino acids

**Supplementary Table 5: Hydrophobicities of amino acids.** Three-letter codes, one-letter codes and hydrophobicities $\varepsilon_i$ of the 20 common amino acid types taken from the 1BPA v1.1 coarse-grained model[8] for DNAJB6b. The hydrophobicity of glutamine (Q) in polyQ is calibrated in this work and set to 0.70.

| Amino acid type | Three-letter code | One-letter code | $\varepsilon_i$ | $q_i$ [$e$] |
|---|---|---|---|---|
| Alanine | ALA | A | 0.7 | 0 |
| Arginine | ARG | R | 0.005 | +1 |
| Asparagine | ASN | N | 0.41 | 0 |
| Aspartatic acid | ASP | D | 0.005 | −1 |
| Cysteine | CYS | C | 0.68 | 0 |
| Glutamine | GLN | Q | 0.33 | 0 |
| Glutamatic acid | GLU | E | 0.005 | −1 |
| Glycine | GLY | G | 0.48 | 0 |
| Histidine | HIS | H | 0.53 | 0 |
| Isoleucine | ILE | I | 0.98 | 0 |
| Leucine | LEU | L | 1 | 0 |
| Lysine | LYS | K | 0.005 | +1 |
| Methionine | MET | M | 0.78 | 0 |
| Phenylalanine | PHE | F | 1 | 0 |
| Proline | PRO | P | 0.65 | 0 |
| Serine | SER | S | 0.45 | 0 |
| Threonine | THR | T | 0.51 | 0 |
| Tryptophan | TRP | W | 0.96 | 0 |
| Tyrosine | TYR | Y | 0.82 | 0 |
| Valine | VAL | V | 0.94 | 0 |
